# Unique genomic features of crAss-like phages, the dominant component of the human gut virome

**DOI:** 10.1101/2020.07.20.212944

**Authors:** Natalya Yutin, Sean Benler, Sergei A. Shmakov, Yuri I. Wolf, Igor Tolstoy, Mike Rayko, Dmitry Antipov, Pavel A. Pevzner, Eugene V. Koonin

## Abstract

CrAssphage is the most abundant virus identified in the human gut virome and the founding member of a large group of bacteriophages that infect bacteria of the phylum Bacteroidetes and have been discovered by metagenomics of both animal-associated and environmental habitats. By analysis of circular contigs from human gut microbiomes, we identified nearly 600 genomes of crAss-like phages. Phylogenetic analysis of conserved genes demonstrates the monophyly of crAss-like phages, which can be expected to become a new order of viruses, and of 5 distinct branches, likely, families within that order. Two of these putative families have not been identified previously. The phages in one of these groups have large genomes (145-192 kilobases) and contain an unprecedented high density of self-splicing introns and inteins. Many crAss-like phages encode suppressor tRNAs that enable readthrough of UGA or UAG stop-codons, mostly, in late phage genes, which could represent a distinct anti-defense strategy. Another putative anti-defense mechanism that might target an unknown defense system in Bacteroidetes inhibiting phage DNA replication involves multiple switches of the phage DNA polymerase type between A and B families. Thus, comparative genomic analysis of the expanded assemblage of crAss-like phages reveals several unusual features of genome architecture and expression as well as phage biology that were not apparent from the previous crAssphage analyses.

## Introduction

During the last few years, the advances of metagenomics have transformed the field of virology by dramatically expanding the virosphere ^1^. The human gut virome is one of the most intensely studied viromes on earth thanks to the obvious health relevance ^2^. However, the majority of the sequences in the gut virome, i.e., the DNA sequences from the virus-like particle fraction, represent virus dark matter, that is, show no significant similarity to any sequences in the current databases ^2–5^. The dark matter can be expected to consist, primarily, of viruses that are, at best, distantly related to the known ones, such that their identification by detecting signature virus protein requires special effort. The most notable case in point is the discovery of crAssphage (after Cross-Assembly), the most abundant human-associated virus. The genome of crAssphage is a circular (or terminally redundant) double-stranded (ds) DNA molecule of approximately 97 kilobase (kb) that was assembled from contigs obtained from mutliple human gut viromes ^6^. The crAssphage is represented in about half of human gut metagenomes and in some of these accounts for up to 90% of the sequencing reads in the virus-like particle fraction. CrAssphage genome sequences have been detected in human gut metagenomes from diverse geographic locations, showing that crAssphage is not only the most abundant virus in some human gut microbiomes but is widely spread across human populations ^2,6–10^.

Initial analysis of the crAssphage genome identified few genes with detectable homologs and failed to establish any relationships with other phages. However, subsequent searches of extended sequence databases using more powerful computational methods resulted in prediction of the functions of the majority of the crAssphage genes and delineation of an expansive group of crAss-like phages in diverse host-associated and environmental viromes ^11^. The gene complements of the crAss-like phages shows a number of distinct features, in particular, a predicted complex transcription machinery centered around an RNA polymerase (RNAP) with an unusual strtucture related to the structure of eukaryotic RNA-dependent RNA polymerases involved in RNA inteference. Recently, the prediction of this unique RNAP has been validated by structural and functional analysis ^12^. A phylogenomic analysis of 250 genomes of crAss-like phages led to the proposal of 4 distinct subfamilies and 10 genera ^13^.

Multiple lines of evidence indicate that the hosts of most if not all crAss-like phages belong to the bacterial phylum Bacteroidetes which represent a dominant component of the human gut microbiome but are, largely, recalcitrant to growth in culture ^6,11,14^. This host range accounts for the fact that the most abundant virus in the human virome remained unknown until the advent of advanced metagenomics, and had been known only as a genome sequence for another 3 years. Nevertheless, recent concerted effort culminated in successful isolation of a crAss-like phage in culture of *Bacteroides intestinalis* ^15^.

Here we report an extensive analysis of large circular contigs from human gut metagenomes that resulted in the identification of nearly 600 diverse genomes of crAss-like phages which are expected to comprise an order of viruses within the class Caudovirecetes, with 5 or 6 constituent families. Some groups of crAss-like phages show unusual genomic features including frequent exchange DNA polymerases of A and B families, extensive utilization of alternative genetic codes in late genes, and unprecedented accumulation of group I self-splicing introns and inteins.

## Results

### Identification of crAss-like phages in human gut metagenomes

In order to obtain a set of complete virus genomes, we extracted all circular contigs (with exact identical regions 50 to 200 basepairs (bp) in length at the ends) from 5,742 human gut metagenomic assemblies resulting in 95,663 contigs (Circular Metagenome Assembled Genomes, hereafter cMAGs). This set of cMAGs presumably should consist of plasmid and virus genomes.

To identify virus cMAGs, we searched the sequences with sequence profiles derived from 421 previously constructed alignments of conserved virus proteins (see Methods). Predicted proteins homologous to one or more of these conserved virus proteins were detected in 4,907 cMAGs. This set of putative virus genomes was then searched with the profile for the large terminase subunit (TerL) of the crAss-like phage group. For the detected TerL homologs, phylogenetic analysis was performed, revealing a major, strongly supported clade that included 596 crAss-like phage cMAGs which formed 169 clusters of distinct genomes sharing less than 90% of similarity at the DNA level. Together with the previously identified related viruses, these genomes comprise the ‘extended group of crAss-like phages’ (Figure 1, Supplementary Table S1). The emerging structure of the group is generally compatible with and extends the findings of the previous phylogenomic analyses ^10,11,13^.

**Figure 1.**
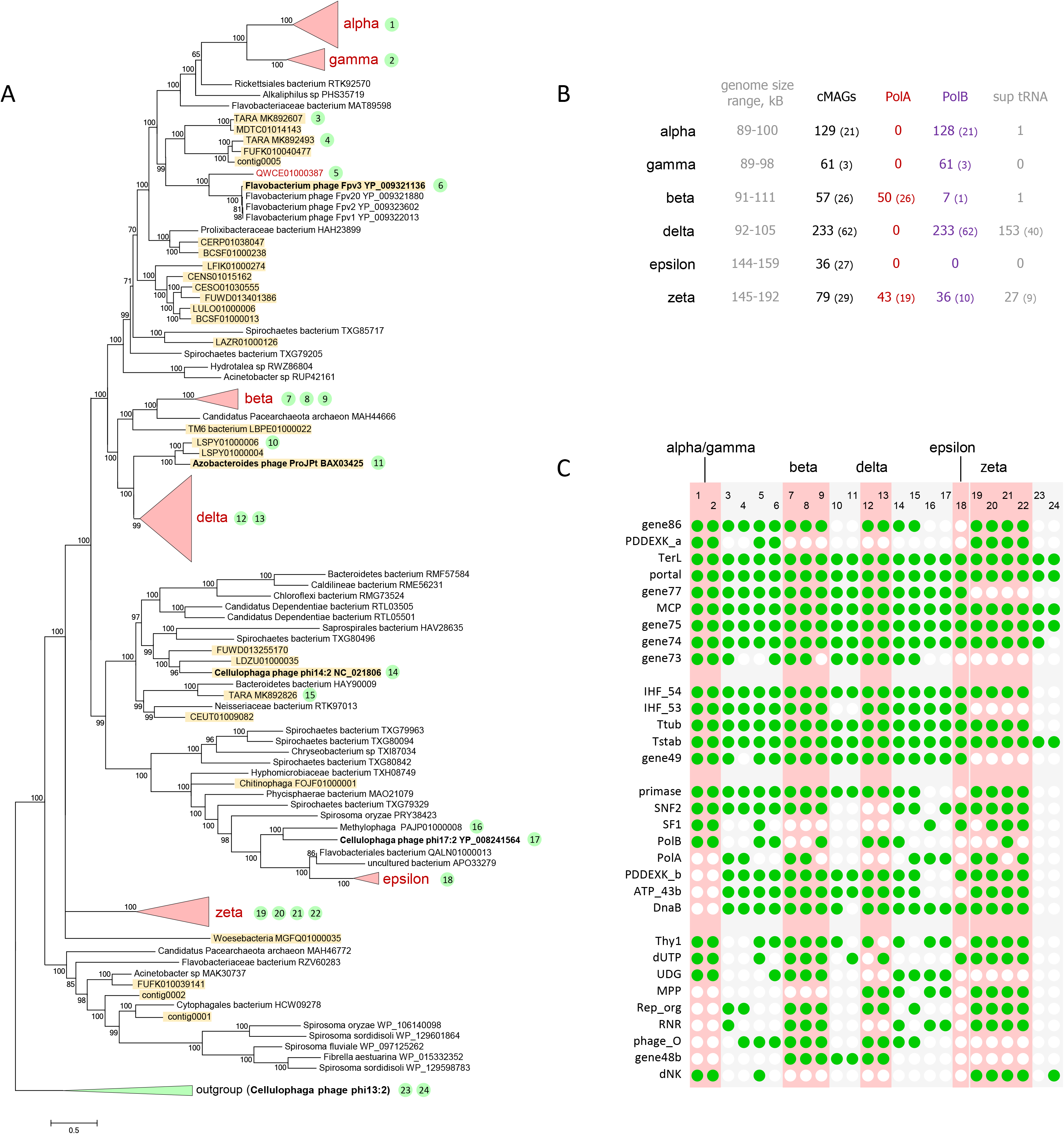
The crAss-like phage assemblage. (A) Phylogenetic tree of the TerL protein of crAss-like phages. Previously analyzed phages are shaded. Numbers in green circles indicate genomes for which the gene conservation pattern is shown in (C). (B) Some key genome features of crass-like cMAGs. In parentheses are numbers on distinct genomes (sharing less than 90% of similarity at the DNA level). (C) Pattern of gene conservation in crAss-like phages. Abbreviations are as follows: TerL, terminase large subunit; portal, portal protein; gene77, putative structural protein (gene 77); MCP, Major capsid protein; gene75, putative structural protein (gene 75); gene74, putative structural protein (gene 74); gene73, putative structural protein (gene 73); IHF_54, IHF subunit (gene 54); IHF_53, IHF subunit (gene 53); Ttub, Tail tubular protein (P22 gp4-like); Tstab, Tail stabilization protein (P22 gp10-like); gene49, Uncharacterized protein (gene 49); gene86, putative structural protein (gene 86); PDDEXK, PD-(D/E)XK family nuclease; DnaB, phage replicative helicase, DnaB family; primase, DnaG family primase; SNF2, SNF2 helicase; SF1, SF1 helicase; ATP_43b, AAA domain ATPase; PolB, DNA polymerase family B; PolA, DNA polymerase family A; Thy1, Thymidylate synthase; dUTP, dUTPase; UDG, Uracil-DNA glycosylase; MPP, metallophosphatase; Rep_Org, replisome organizer protein; RNR, ribonucleotide reductase; phage_O, Bacteriophage replication protein O (gene10 of IAS virus); gene48b, phage endonuclease I; dNK, deoxynucleotide monophosphate kinase.

We further sought to characterize the host range of crAss-like phages, and to that end, performed a search for CRISPR spacers matching the genome of these phages^16^ (see Methods for details). This approach identified potential hosts for 466 of the 673 crAss-like phages. An overwhelming majority of the spacers with reliable matches came from CRISPR arrays in the genomes of different subdivisions of the phylum Bacteroidetes (Supplementary Table S1), supporting and expanding the previous host assignments. Nevertheless, several reliable spacer matches were detected in CRISPR arrays from bacterial phyla other the Bacteroidetes (Supplementary Table S1).

### Conserved and group-specific genomic features in the extended crAss-like phage assemblage

Examination of the TerL tree suggests a coarse-grain structure of the relationships among the crAss-like phages, which include the 596-strong human gut ‘crAss-virome’, the previously analyzed environmental contigs and several independently isolated phages. The monophyly of the previously proposed alpha-gamma, beta, and delta subfamilies of crAss-like phages was confirmed; in addition, two previously unknown groups were identified, dubbed here zeta and epsilon (Figure 1 and Supplementary Table S1). Zeta and epsilon groups comprise the deepest branches in the TerL tree outgrouped by environmental phages that are not considered parts of the crAss-like assemblage. The 5 groups, alpha-gamma, beta, delta, epsilon and zeta groups account for 595 of the 596 cMAGs in the human gut crAss-virome. One cMAG did not belong to any of these 5 clades but rather grouped with the Flavobacteroides phage Fpv3 (a phage infecting a fish pathogenic bacterium); this might not be a native human gut phage (Figure 1A).

The alpha-gamma group is best characterized and annotated, and includes the ‘crAssphage clade’ (alpha) where the original crAssphage belongs ^10,11^. In our initial analysis ^11^, the beta group was represented by a single complete genome, immune deficit-associated (IAS) phage; since then, this group has been expanded to include a variety of human gut metagenomic contigs^13^ along with ΦCrAss001, the only cultured crAss-like phage ^15^. The delta group was initially represented by several incomplete phage genomes ^11^ and was subsequently substantially expanded ^13^. The delta group is currently the largest one in the gut crAss-virome (Figure 1B). The epsilon group is dominated by gut cMAGs but additionally includes *Cellulophaga* phage phi17:2 and two GenBank contigs, *Methylophaga* PAJP01000008 and Flavobacteriales bacterium QALN01000013. The phage genomes in the epsilon group are about 150 kb long, i. e. about 50% larger than the original crAssphage genome. The zeta group consists entirely of MAGs from the human gut microbiomes analyzed here and some previously reported human gut and oral metagenome contigs ^17^, with no homologs, detectable by BLASTP search, among known phages or any sequences in GenBank. These phages show considerable within-group diversity and have genomes in the range of 145-192 kb, some of which are nearly twice as large as the original crAss-like genomes.

The genomes of crAss-like phages contain 3 readily discernible blocks of genes that encode: 1) virion components and proteins involved in particle assembly, 2) components of the replication apparatus, 3) components of the transcription machinery (Figure 2). The structural block is overall the most highly conserved one and includes the 8 genes that are shared by all crAss-like phages: MCP, TerL, portal protein, Integration Host Factor (IHF), tail stabilization protein, tail tubular protein, and uncharacterized genes 74 (beta and delta groups encompass tandem duplications of this gene) and 75; in addition, gene 86 that is adjacent to *terL* encodes a small protein that is likely to represent the terminase small subunit despite the lack of detectable similarity TerS of other phages (the genes are numbered according to the crAssphage genome annotation). The presence of the IHF, which is likely to contribute to DNA compactification during virion assembly, and of the two conserved uncharacterized genes in the structural block can be considered the genomic signature of the crAss-like genes. Three more genes including a paralog of IHF are present in (nearly) all crAss-like genes except for the zeta group, and one more gene is missing in both in the zeta and epsilon groups (Figure 1C).

**Figure 2.**
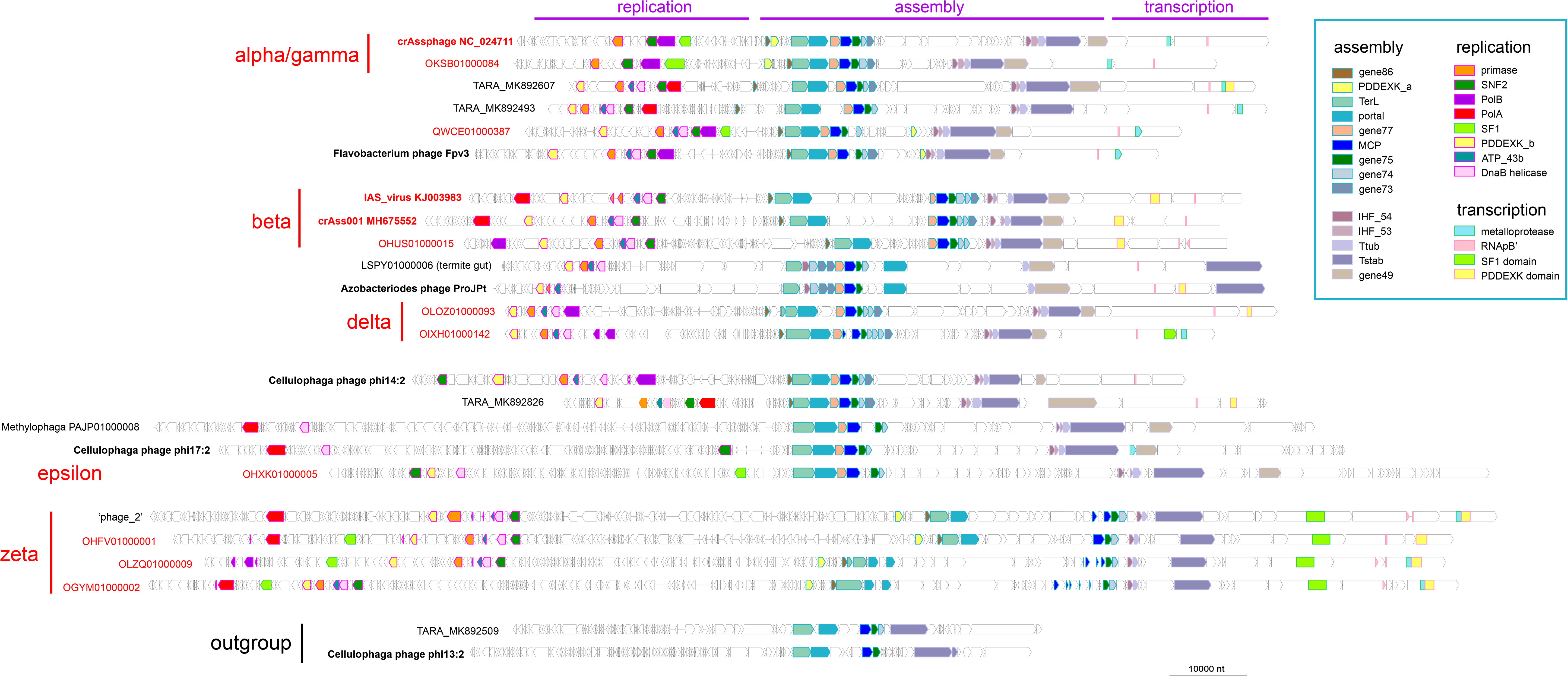
Genome organizations of representative crAss-like phages. The circular genomes were linearized by breaking the circle after the transcription gene block. Conserved proteins are abbreviated as in Fig 1C; RNApB’ marks the conserved DxDxD motif of the large RNAP subunit homologous to the bacterial β’ subunit.

In the replication block, the four highly conserved genes are DnaG family primase, two helicases of DnaB and SNF2 families, and an AAA+ superfamily ATPase. Each of these genes is represented in the majority of crAss-like phages including the zeta group which implies that they were present in the last common ancestor of these phages but were differentially lost in the alpha-gamma, delta or epsilon group (Figure 1C, Supplementary Figure 1). In accord with previous observations, phylogenetic analysis of the primase from the extended set of crAss-like phages demonstrates the single origin of the phage primase from DnaG of Bacteroidetes (Supplementary Figure 2). With the exception of the epsilon group and a few phages from environments other than human gut, the crAss-like phages encode a DNA polymerase (DNAP) of either A or B family (PolA and PolB, respectively). The evolution of the DNAPs of crAss-like phages is discussed in detail in the next section. In addition to the proteins directly involved in replication, crAss-like phages encode several enzymes of nucleotide metabolism, in particular, thymidylate synthase (Thy1), dUTPase, Uracil-DNA glycosylase (UDG) and ribonucleotide reductase (RNR). In this group of enzymes, Thy1, dUTPase and RNR appear to be ancestral among the crAss-like phages, albeit lost in some members (Figure 1C). As this analysis shows, some of the crAss-like phages, particularly, those in the zeta group that have the largest genomes in the entire assemblage, encode a complex replication apparatus complemented by enzymes of deoxynucleotide metabolism. By contrast, phages in the epsilon group possess a minimalist replication apparatus.

The transcription gene block that is present in all crAss-like phage genomes, except for the epsilon group, consists of several large, multidomain proteins, one of which contains the DxDxD motif that is conserved in one of the large subunits of bacterial, archaeal and eukaryotic RNAPs and is an essential part of the catalytic site (Figure 3). The predicted RNAPs of crAss-like phages are extremely divergent in sequence from all known RNAPs and from each other, suggestive of unusually high evolutionary rates. The recent structural and functional study of the RNAP of *Cellulophaga baltica* phage phi 14:2 of confirms the involvement of this large protein in the transcription of the phage early genes and shows that it contains two double-psi beta-barrel (DPBB) domains within a single polypeptide, unlike cellular RNAPs in which the two DPBB domains belong to the two largest subunits ^12^.

**Figure 3.**
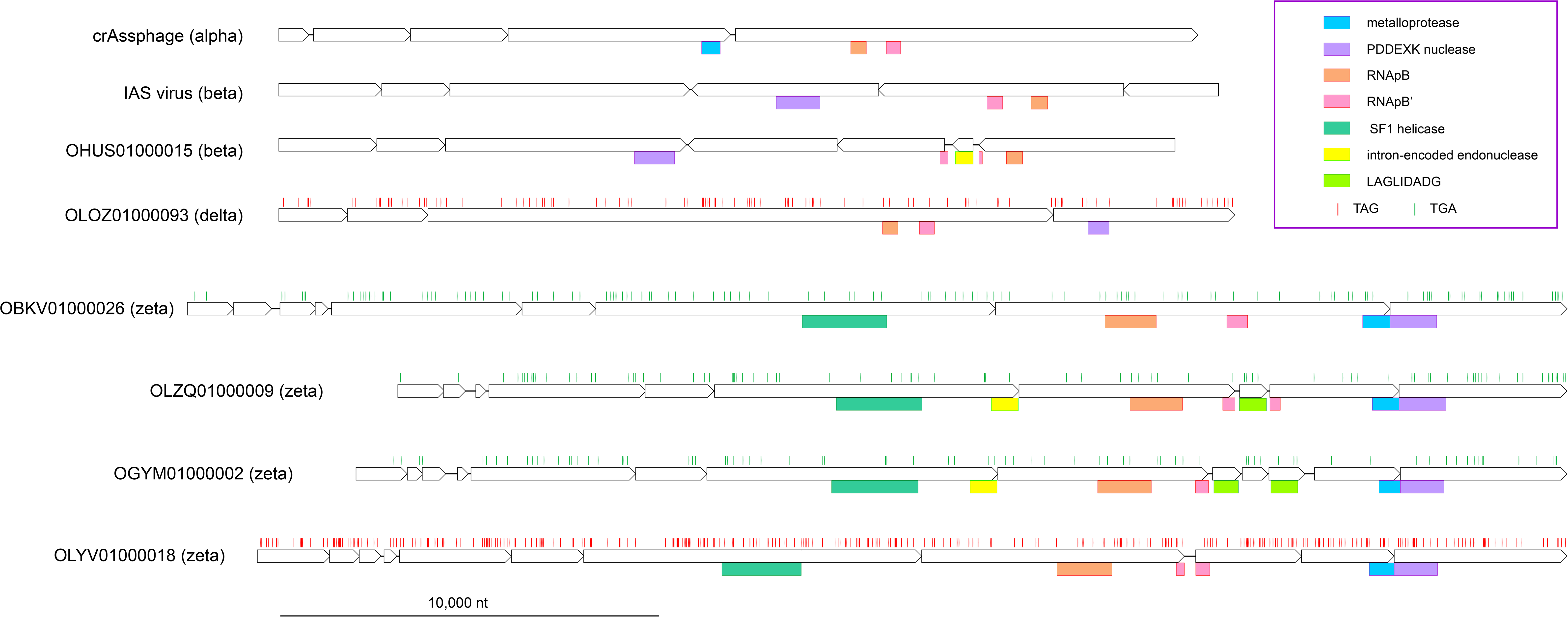
Transcription gene block in crAss-like phages. The predicted genes are shown by empty block arrows, and recognized conserved domains are indicated by colored rectangles. Vertical bars indicate in-frame stop codons. LAGLIDADG is intron-encoded endonuclease-maturase.

A distinct feature of the crAss-like phages (with the exception of some members of the epsilon group) is the presence of at least one nuclease of the PDDEXK family, whereas many phages encode 2 or even 3 such nucleases (Figure 2). Apparently, these nucleases perform more than one function in phage reproduction because one variety is encoded within the structural block (PDDEXK_a), and another one within the replication block (PDDEXK_b), whereas some phages contain a PDDEXK domain in one of the large proteins associated with transcription (Figure 2, 3).

#### Switching of DNA polymerases in crAss-like phages

With the exception of the epsilon group, crAss-like phages contain a gene within the replication block that encodes a DNA polymerase of either A family or B family (the two families contain distantly related variants of the core Palm domain^18^) (Figure 1B). In some of the groups of crAss-like phages, one or the other DNA polymerase is used exclusively or preferentially, for example, PolB in the alpha-gamma group, and PolA in the beta group, but in other groups, in particular, zeta, PolA and PolB are mixed on a much finer scale (Figure 4, Supplementary Figure 3). The location of the DNAP gene is mostly consistent within a group but can differ between the groups, for example, upstream of the primase gene in alpha, and downstream of the primase in beta (Supplementary Figure 1). Notably, in the groups where the two DNAP varieties are present alternatively, they typically occupy the same position, that is, appear to be replaced *in situ* (Supplementary Figure 4).

**Figure 4.**
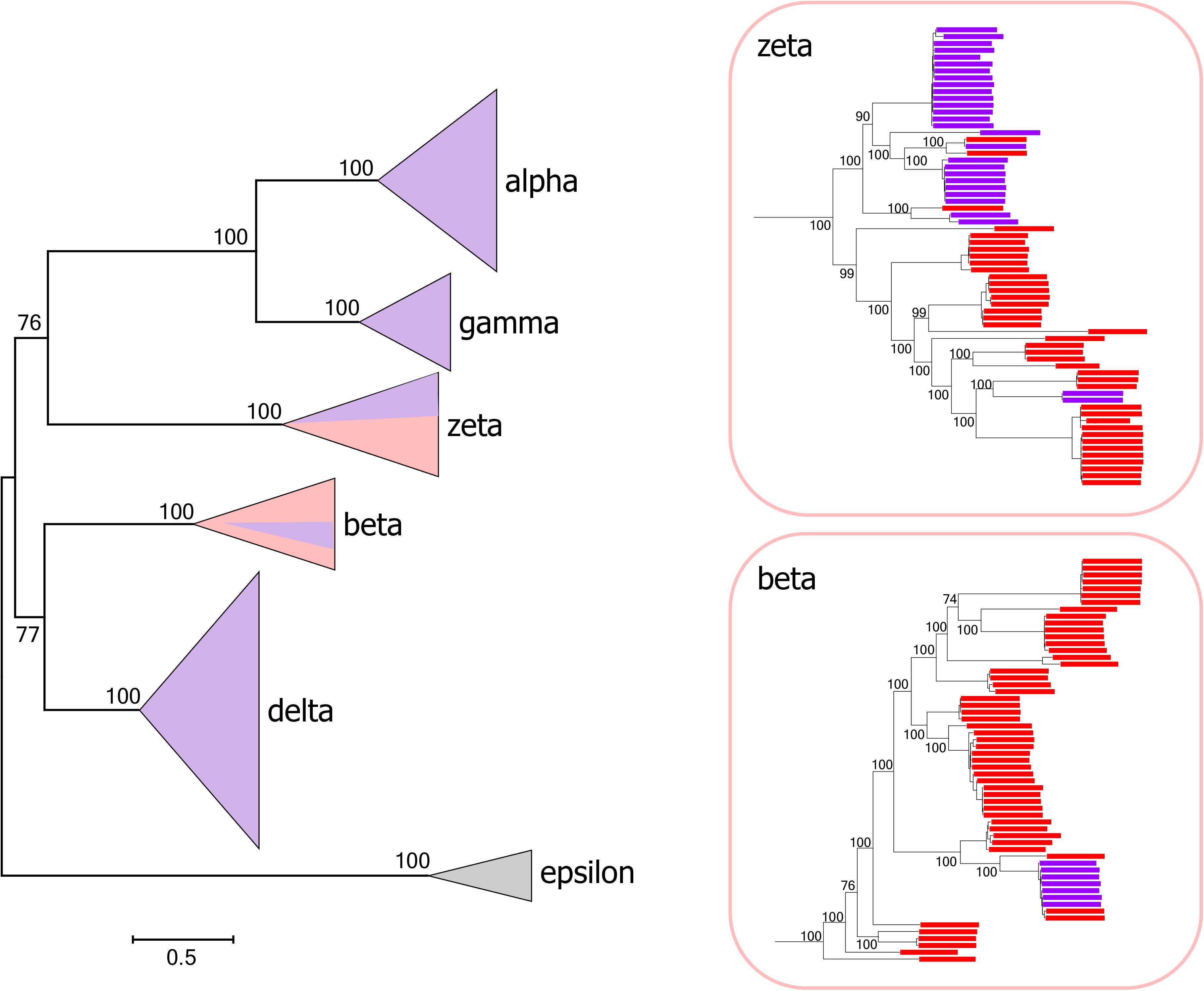
DNA polymerase switches in the evolution of crAss-like phages. The left panel shows a schematic phylogenetic tree of TerL for the entire crAss-like phage assemblage, with the branches collapsed into triangles. The right panels show the expanded beta and zeta branches. The portions of the triangles (left panel) and the tree leaves (right panels) are colored by the DNAP type, red for PolA and purple for PolB.

Because none of the extant crAss-like phages encode two DNAPs, it is natural to assume that their last common ancestor also possessed only one DNAP. The extant distribution of PolA and PolB, together with the phylogenetic analysis of both DNAP families, imply numerous, independent events of both xenologous gene displacement (by a DNAP gene of same family but from a distant clade) and non-orthologous gene displacement (PolA with PolB or vice versa), often occurring *in situ*, that is, without disruption of the gene organization in the phage genomes. Thus, parsimonious reconstruction of the ancestral state does not seem to be feasible, so that both PolA and PolB are equally suitable candidates for the role of the ancestral DNAP. Phylogenetic analysis of both PolA and PolB indicate their monophyly among the crAss-like phages except, possibly, for some distant variants in the zeta group, so that displacement appears to have involved, primarily, among the crAss-like phages (Supplementary Figure 3). The presence of different DNAPs in equivalent positions in many pairs of closely related phages implies that DNAP gene displacement occurs by homologous recombination. Given that the 5 groups of crAss-like phages remain, to a large extent, monophyletic in the DNAP phylogenies (Supplementary Figure 3), it seems likely that, within the groups, displacement occurs, primarily, by homologous recombination between closely related genomes during coinfection, followed by sequence divergence. However, the observed mixing between groups in the DNAP phylogenies, for example, the presence of multiple members of the zeta group within the beta branch of PolB, implies gene displacement between distantly related phages that require mechanisms other than homologous recombination (Supplementary Figure 5).

#### Alternative coding strategies in crAss-like phages

Typically, annotation of a bacterial virus genome starts with predicting protein-coding genes (Open Reading Frames, ORFs) and producing the conceptual translation of the ORFs into protein sequences ^19^. When attempting such annotation of the genomes of the crAss-like assemblage, we encountered several phenomena that substantially impeded the analysis. Many of the genomes contain short ORFs encoding fragments of conserved proteins. Some of these fragments lie in the same frame and are interrupted only by standard stop codons; others are also in the same frame but are separated by longer in-frame inserts; yet others are in different frames and are separated by nucleotide sequences that are rich in stop codons in all co-directed frames. Even more unusually, some of the crAss-like phage genomes, particularly, those of the zeta group, contain regions that do not encode any recognizable proteins but rather are occupied by anomalously short ORFs of different polarities. Many of the predicted protein sequences in these regions do not match even when the rest of the phage genomes are closely similar.

Some of the crAss-like phage genomes (for example, Eld241-t0_s_1 of the delta group; Supplementary Figure 6) contain almost no long ORFs when translated under the standard genetic code with three standard stop codons. However, these genomes contain many ORFs encoding homologs of known phage proteins including TerL, portal, and RNAP subunits when translated with an alternative code in which TAG encodes an amino acid (most likely, glutamine) instead of being a stop codon. Thus, these phages apparently use such an alternative genetic code; indeed, the entire genome of Eld241-t0_s_1 has been translated using the code with TAG reassigned for glutamine ^13^.

Alternative genetic codes in phages including extensive opal (TGA) and amber (TAG) stop codon reassignments have been reported previously, especially, in human-associated metagenomes ^17,20,21^. We used the presence of standard stop codons in several readily identifiable, nearly ubiquitous genes (TerL, MCP, portal) as evidence of likely stop codon reassignment, and the contigs with clear evidence of alternative code use were translated using the appropriate code tables (Supplementary Figure 7, Supplementary Table S1). Altogether, we identified 243 crAss-like phages, mostly, from the beta, delta and zeta groups, using alternative codes, with either TAG reassigned for glutamine or TGA reassigned for tryptophan (Supplementary Table S1). In most cases, stop codon reassignment was observed only in parts of the genome that encompass putative late genes. CrAss-like phages apparently can switch genetic codes at short phylogenetic distances; for example, different branches within the zeta group reassign either TAG or TGA, or use the standard code (Supplementary Figure 5).

One of the common mechanisms for codon reassignment employs suppressor tRNAs that have an anticodon complementary to one of the standard stop codons and are charged with an amino acid. Many genomes of crAss-like phages, especially, in the beta and zeta groups, encode multiple tRNAs including putative suppressors (Supplementary Figure 8, Supplementary Table S1). However, the evidence of codon reassignment is not in a prefect agreement with the presence of suppressor tRNAs (as exemplified for the zeta group in Supplementary Figure 5), suggesting alternative mechanisms for genetic code fine-tuning in these phages.

We traced several cases of suppressor tRNA emergence. In one case, a suppressor tRNA emerges in one a phage genome via a single point mutation in the anticodon of a tRNA (TTG^Gln^→TTA^Sup^) (Supplementary Figure 9). The block of three tRNAs (Tyr, Ley, Gln) is present in several closely related genomes from the alpha group, and one of these (OKSC01000115) carries the mutation that transforms tRNA^Gln^ into a TAA stop codon suppressor. This genome as well as those of its close relatives in the alpha group show no traces of the TAA codon (or any other stop codon) reassignment. This is an apparent case of suppressor pre-emergence that would open the path for stop codon reassignment in the descendants of the mutant phage and could be a general mechanism of code switch evolution. Indeed, OFRY01000050 genome from the beta group acquired a TAG suppressor tRNA, presumably, from a bacterial source (Supplementary Figure 10). Unlike its closest relatives, this genome started to accumulate TAG codons in conserved genes, such as TerL, in an apparent case of ongoing progression along the alternative coding path.

#### Unprecedented infestation of crAss-like phage genomes with self-splicing introns and inteins

In many bacteriophages, some genes are interrupted by Group I or Group II self-splicing introns and/or inteins ^22^. However, some of the crAss-like phages are characterized by unprecedented density of introns and inteins. The genomes in the alpha-gamma group that have been primarily studied previously lack MGE insertions which enabled straightforward annotation. In contrast, phages of the delta and, especially, zeta groups, harbor numerous introns and inteins (Supplementary Figure 11). In phages of the delta group, group I introns are typically inserted in the core phage genes including MCP, PolB, primase, and ATP_43b, whereas TerL typically contains an intein. The phage genomes in the zeta group contain even more introns and inteins inserted in many different genes (Supplementary Figure 11). Together with the use of alternative genetic codes, the massive infestation of these genomes with introns and inteins severely complicates annotation.

An example of an intron-rich gene is the MCP gene of the zeta group cMAG OLRF01000054 (Figure 5A). The MCP ORF is split into 5 fragments across two co-directed coding frames; three of the four segments between the ORF fragments encode domains typical of Group I self-splicing introns. Within the recognizable protein coding sequences, no standard stop codons are present, so there is apparently no alternative coding in this gene. In the OBYQ01000140 cMAG (delta group), TerL is encoded in three in-frame fragments that are separated by two inteins of the Hint-Hop type. Both the TerL and the intein parts of the ORF contain multiple TAG codons indicative of the use of an alternative code (Figure 5B). Thus, in this case, intein insertion and alternative coding are combined within the same gene. PolB of the zeta group cMAG OLWB01000021 is encoded in four fragments across three frames; two of the regions separating the coding sequences encode intron-specific domains, and both the PolB-coding and intron ORFs contain multiple TGA codons that have to be suppressed during translation (Figure 5C).

**Figure 5.**
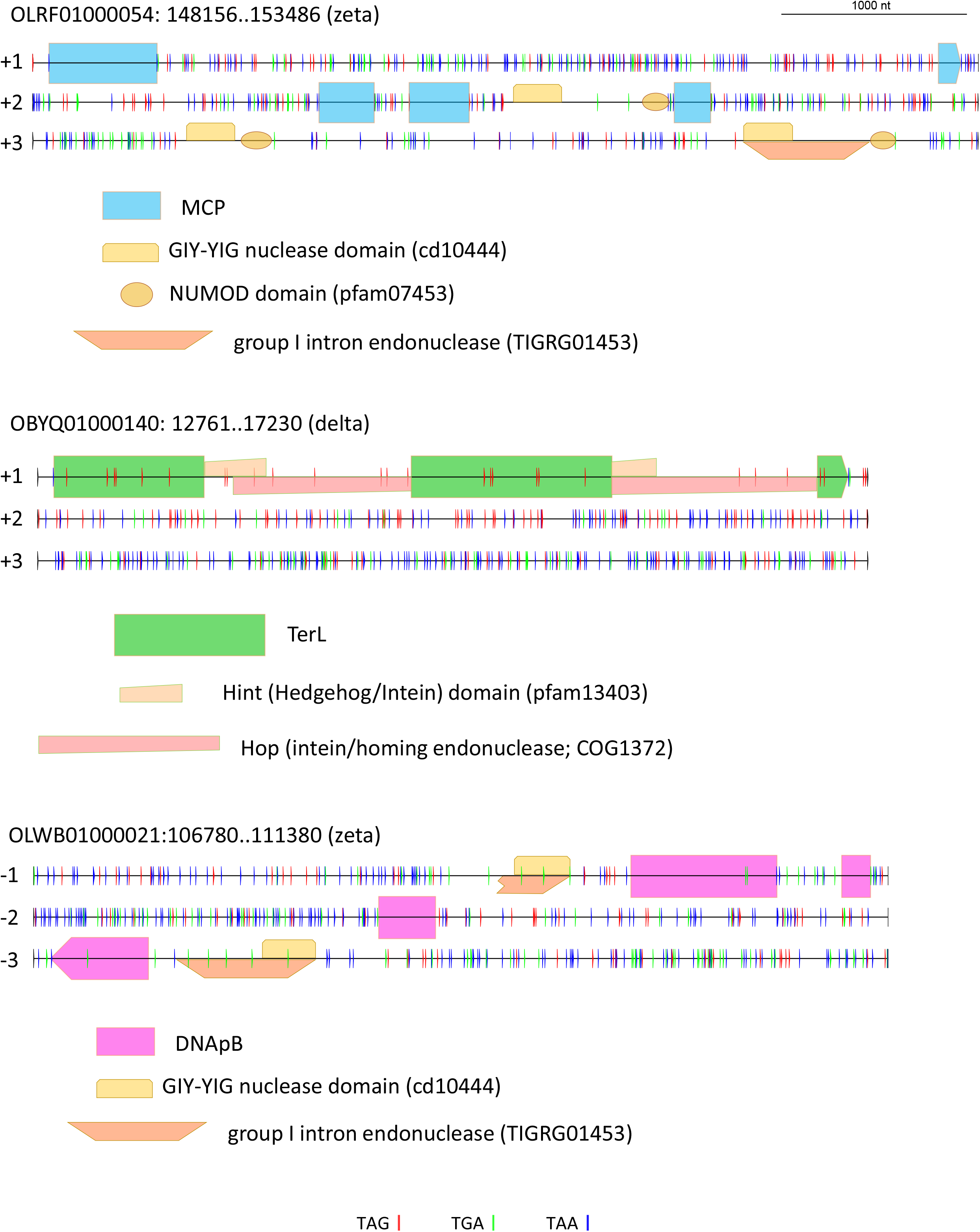
Examples of stop codon reassignments, intein and self-splicing intron insertions in conserved genes of crAss-like phages. Horizontal lines denote co-directed reading frames in the nucleotide sequence; short vertical bars indicate standard stop codons (red: TAG, green: TGA, blue: TAA); colored shapes indicate domains or domain fragments, mapped to the nucleotide sequence.

In the zeta group, the ORFs coding for the predicted RNAP subunits are even more drastically disrupted, with numerous stop codons and frameshifts some of which occur within highly conserved motifs, such as the DxDxD motif of RNAPB’ (Figure 3, Supplementary Figure 12). The predicted protein boundaries vary considerably within clades of closely related phages. It seems likely that these phages, in addition to RNA and protein splicing, and stop codon reassignment, employ additional, yet unknown expression mechanisms.

## Discussion

The identification and genome analysis of the crAss-like phages described here substantially expand this group of viruses and confirm its prominence in the human gut virome where the crAss-virome accounts for about 12% of the diversity of the circular virus genomes. CrAss-like phages can be readily defined as a group and differentiated from other phages through the phylogenetic coherence of the conserved genes as well as distinct gene signatures, such as unique genes in the structural module and the giant proteins comprising the transcription apparatus. Within the CrAss-like assemblage, however, there is extensive diversity, in terms of the genome size and content as well as expression strategy. The originally discovered crAssphage and its close relatives^6,10,11,13^ remain the most abundant viruses in the human-associated virome, but they are relatively small compared to phages in other groups, lack some characteristic genes, and therefore, cannot serve as typical representatives of the crAss virome diversity. The crAss-like phages appear ripe to be formally placed into the framework of virus taxonomy. In the recently adopted multi-rank taxonomic structure, crAss-like phages can be expected to become an order within the existing class *Caudovirecetes* (tailed viruses of bacteria and archaea), with alpha-gamma (or, possibly, alpha and gamma separately), beta, delta, epsilon, and zeta groups becoming families within this order.

CrAssphage and other members of the alpha-gamma group described previously have comparatively simple genome organizations and, apart from the unique transcription machinery, employ conventional strategies for genome expression. This historical accident greatly facilitated the initial genome annotation and analysis. However, this is not the case for the much broader assemblage of crAss-like phages analyzed here. In particular, many phages in the Beta, Delta and Zeta groups employ non-standard genetic codes, and those in the latter group, have an unprecedented high density of introns and inteins. Recoding enabled by the capture and adaptation of the suppressor tRNA can be perceived as an anti-defense strategy to impair the production of host proteins, including those involved in defense, by stop-codon readthrough, while faithfully translating the phage proteins. Whether the extreme accumulation of introns and inteins, along with potential additional, unknown forms of gene disruption, in the Zeta group phages is some type of adaptive strategy or a rampage of MGE that spun out of control due to some unknown facet of the lifestyle of these phages, remains unclear.

The frequent DNAP type switches across the entire crAss-like phage assemblage represent an enigmatic phenomenon that, to our knowledge, has not been observed previously. The pervasiveness of these switches implies a strong selective pressure that might have to do with an uncharacterized, distinct defense mechanism characteristic of Bacteroidetes. Perhaps, such defense might involve DNAP inhibition so that DNAP type switch becomes a means of escape. The high prevalence of *in situ* DNAP replacement suggests that co-regulation of genes involved in replication is important for the reproduction of crAss-like phages.

The phylogenomic study of crAss-like phages not only advances our knowledge of the human gut virome, but reveals fascinating, poorly understood aspects of phage biology. In particular, the alternative coding strategies employed by prokaryotic viruses remain to be explored in full. Along with other recent discoveries, such as the plethora of megaphages encoding enormously rich protein repertoires ^20,21^, these findings show that, although phages have been classic models of molecular genetics for 8 decades, there is more to be learned about them than we already know.

## Methods

### Identification of cMAGs in the human gut virome

5,742 whole-community metagenome assemblies generated from human fecal samples were downloaded from the NCBI Assembly database ^23^ (accessed 8/2019). To limit the analyzed dataset to complete, fully assembled genomes, 95,663 ‘circular’ contigs (50-200 bp direct overlap at contig ends) were extracted from the assemblies.

We used 421 phage-specific protein alignments, 117 custom ones and 304 from the CDD database ^24^, and created a set of the corresponding HMMs using hhmake ^25^. Proteins in the 95,663 contigs were predicted using Prodigal ^26^ in the metagenomic mode, and searched against the set of 421 phage-specific HMMs using hhsearch ^25^, with the relaxed e-value cut-off <0.05. 4,907 contigs with at least one hit were selected for subsequent analysis.

### Clustering of virus genomes

The set of 673 extended crAss-like genome assemblies was searched against itself using MEGABLAST ^27^ with no low-complexity filtering, e-value threshold of 10^−8^ and the identity threshold for the highest-scoring hit of 90%. Pairs of genomes where the coverage of the query genome by hits was at least 90% were linked; connected clusters were extracted from the linkage graph. Altogether, the set formed 221 cluster; the 596 crAss-like cMAGs from the gut metagenomes belong to 169 of these clusters.

### Identification of potential hosts of the crAss-like phages

258,077 bacterial and 4,975 archaeal assemblies in the NCBI Genome database ^23^ were scanned for CRISPR arrays using CRISPRCasTyper ^28^; this procedure yielded 3,455,966 CRISPR spacers in 236,854 arrays from 99,448 individual assemblies. A separate curated CRISPR spacers database, derived from high-quality completely sequenced genomes, containing 274,663 spacers ^29^, was used in parallel.

The results of the BLASTN search of 673 crAss family assemblies against the spacer databases were analyzed as follows: a link between a virus assembly and a contig (genome partition) bearing a CRISPR array was established when BLASTN yielded one or more hits with at least 90% of the spacer positions matching the corresponding phage sequence or two or more hits with at least 80% identity. Then, the source organism of the contig with the highest-scoring hit was selected as the potential host, with the preference given to spacers derived from completely sequenced genomes.

The taxonomic assignment of non-Bacteroidetes potential host contigs was verified by running a translating search against the protein database derived from completely sequenced genomes; the lowest-level taxa, accounting for at least 75% of all hits were considered a ‘safe’ taxonomic affiliation (e.g. JUJT01000004.1, deposited in GenBank as *Pectobacterium brasiliense*, produced best hits into complete genomes of Enterobacteriales, confirming this order as the most probable taxonomic assignment for this contig).

### Gene calling with alternative genetic codes and tRNA scan

A modified version of the Prodigal program ^26^, which allows assigning amino acids to codons that serve as stop codons in the standard bacterial genetic code (-g 11), was used to translate the genes with suspected stop codon reassignment. All contigs were searched for the presence of tRNAs using tRNA-scan-SE (v. 2.0) ^30^ employing a bacterial model of tRNAs (-B) and bitscore cutoff of 35.

### Protein sequence analysis

A combination of PSI-BLAST ^31^ searches using the CDD database ^24^ profiles and custom profiles as queries, and HHPRED ^25^ searches was used for domain identification and functional annotation of proteins.

Multiple sequence alignments were constructed using MUSCLE ^32^. Phylogenetic reconstruction was performed using the IQ-TREE program, with the evolutionary models selected by IQ-TREE ^33^.

Detection of cMAG homologs was performing by running BLASTP search of predicted proteins against the subset of proteins in the NCBI NR database with taxonomic label “Viruses”; e-value threshold of 0.0001; no low complexity filtering.

## Data availability

Genome and protein sequences analyzed in this work, functional annotations, custom profiles, multiple sequence alignments, and phylogenetic trees are available from: ftp://ftp.ncbi.nih.gov/pub/yutinn/crassfamily_2020/

## Supplementary figures

**Suppl Figure 1** Group-specific genome maps for replication gene block

**Suppl Figure 2** Phylogenetic tree of DnaG family primase

**Suppl Figure 3** Phylogenetic trees of PolA and PolB

**Suppl Figure 4** Two examples of in situ DNAP replacement

**Suppl Figure 5** – Genomic features of zeta group phages (DNAP type, recoding, Sup tRNAs detected) mapped on a TerL phylogenetic tree.

**Suppl Figure 6** TAG, TGA, and TAA codons across the genomes of crAssphage and contig Eld241-t0_s_1 (of Guerin ea 2018)

**Suppl Figure 7** Three zeta group phage genomes, OLOE01000004 (A), OKSX01000010 (B), and OIXA01000019 (C), translated with standard bacterial code (top row), standard bacterial code with TAG read-trough (middle row), and standard bacterial code with TGA read-trough (bottom row). Accounting for read-trough TAG and TGA result in longer orfs and higher coding density in parts of OKSX01000010 and OIXA01000019 genomes.

**Suppl Figure 8** Genome size, number of tRNAs and suppressor tRNAs in crass-like cMAGs compared to other gut phage cMAGs (4-300 kb-long).

**Suppl Figure 9** Emergence of suppressor tRNA. A, TerL phylogenetic tree of alpha/gamma group phages. B, a fragment of multiple sequence alignment of tRNA genes of six closely related genomes.

**Suppl Figure 10** A case of suppressor tRNA within beta group. A, TerL phylogenetic tree of beta group phages. B, A fragment of TerL gene alignment; the first three TAGs reassigned to Q in OFRY01000050 genome are shown; the whole TerL gene of OFRY01000050 has 15 reassigned TAG codons. C, A dawn of alternative coding in OFRY01000050 genome. Four closely related genomes were translated with standard bacterial code and standard bacterial code with TAG read-trough. Regions of alternative coding in OFRY01000050 are marked with dashed boxes.

**Suppl Figure 11** Abundance of introns and intein domains in crass-like phages. Whole genome maps for representative alpha/beta, delta, zeta, and epsilon phages are shown; whole-genome annotations of these genomes are here: ftp://ftp.ncbi.nih.gov/pub/yutinn/crassfamily_2020/

**Suppl Figure 12** Structure of the transcription gene in zeta group phages. Conserved domains were identified with translating blast searches against a custom set of profiles and superimposed on genome map constructed with Prodigal.

