## Supplementary figures for "Unique genomic features of crAss-like phages, the dominant component of the human gut virome"

### Supplementary Figure 1

A

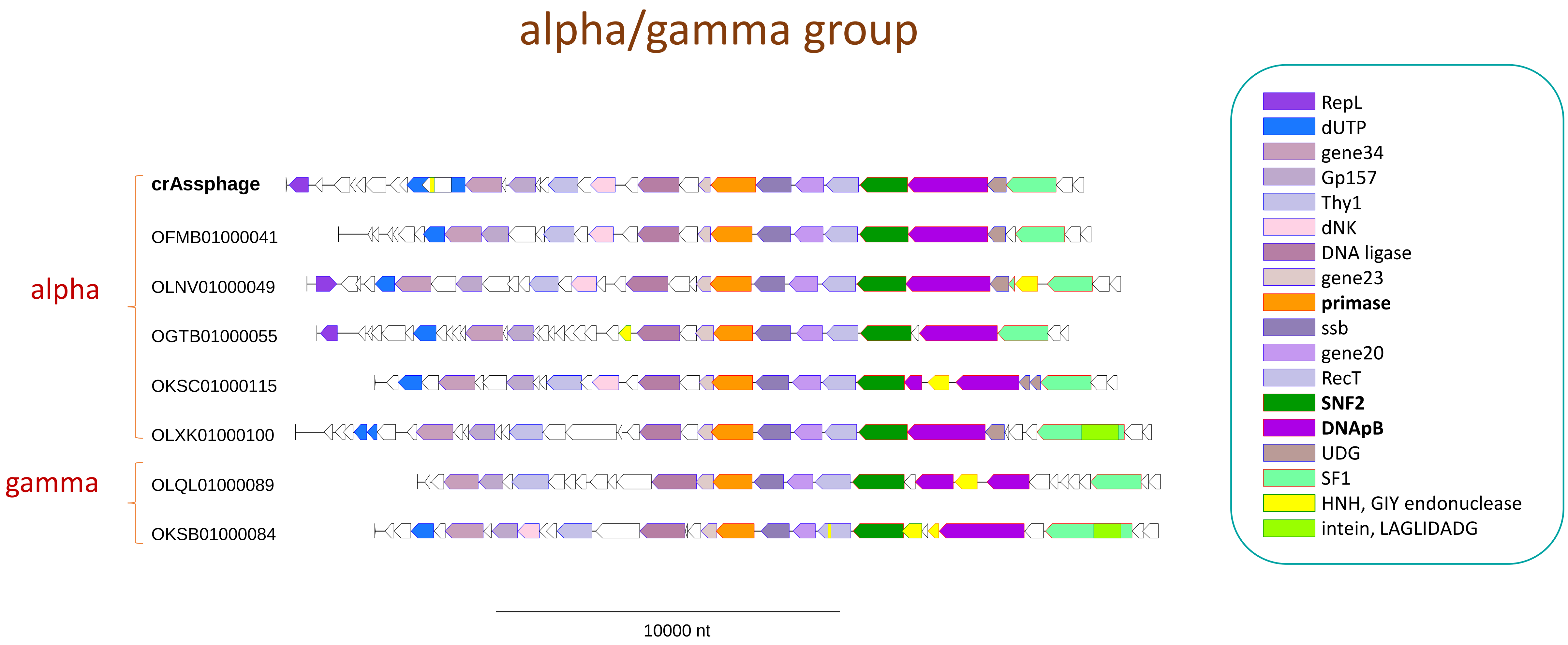

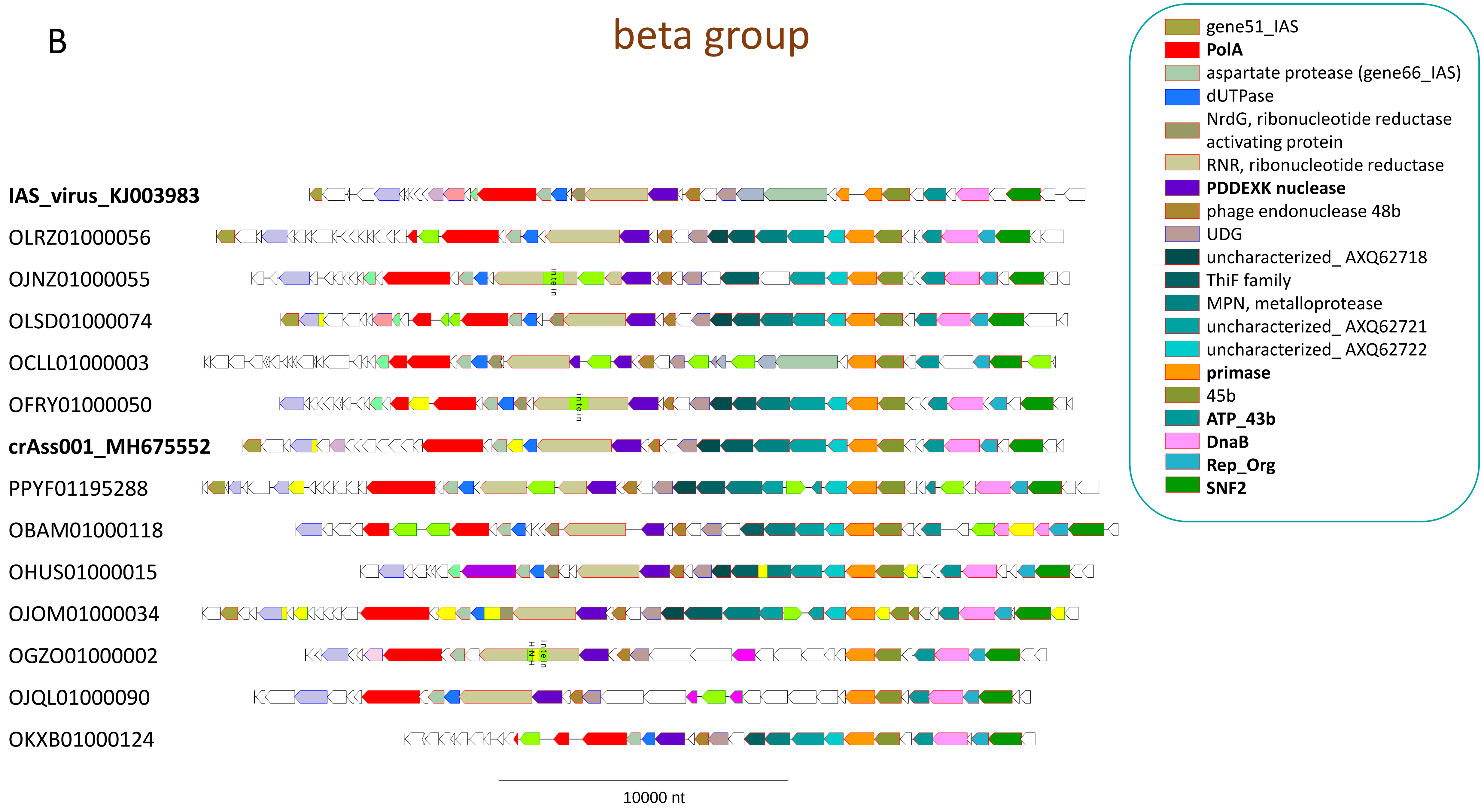
B
beta group
IAS_virus_KJ003983
OLRZ01000056
OJNZ01000055
OLSD01000074
OCLL01000003
OFRY01000050
crAss001_MH675552
PPYF01195288
OBAM01000118
OHUS01000015
OJOM01000034
OGZO01000002
OJQL01000090
OKXB01000124
intein
HNH
10000 nt
gene51_IAS
PolA
aspartate protease (gene66_IAS)
dUTPase
NrdG, ribonucleotide reductase activating protein
RNR, ribonucleotide reductase
PDDEXK nuclease
phage endonuclease 48b
UDG
uncharacterized_ AXQ62718
ThiF family
MPN, metalloprotease
uncharacterized_ AXQ62721
uncharacterized_ AXQ62722
primase
45b
ATP_43b
DnaB
Rep_Org
SNF2

c

### delta group

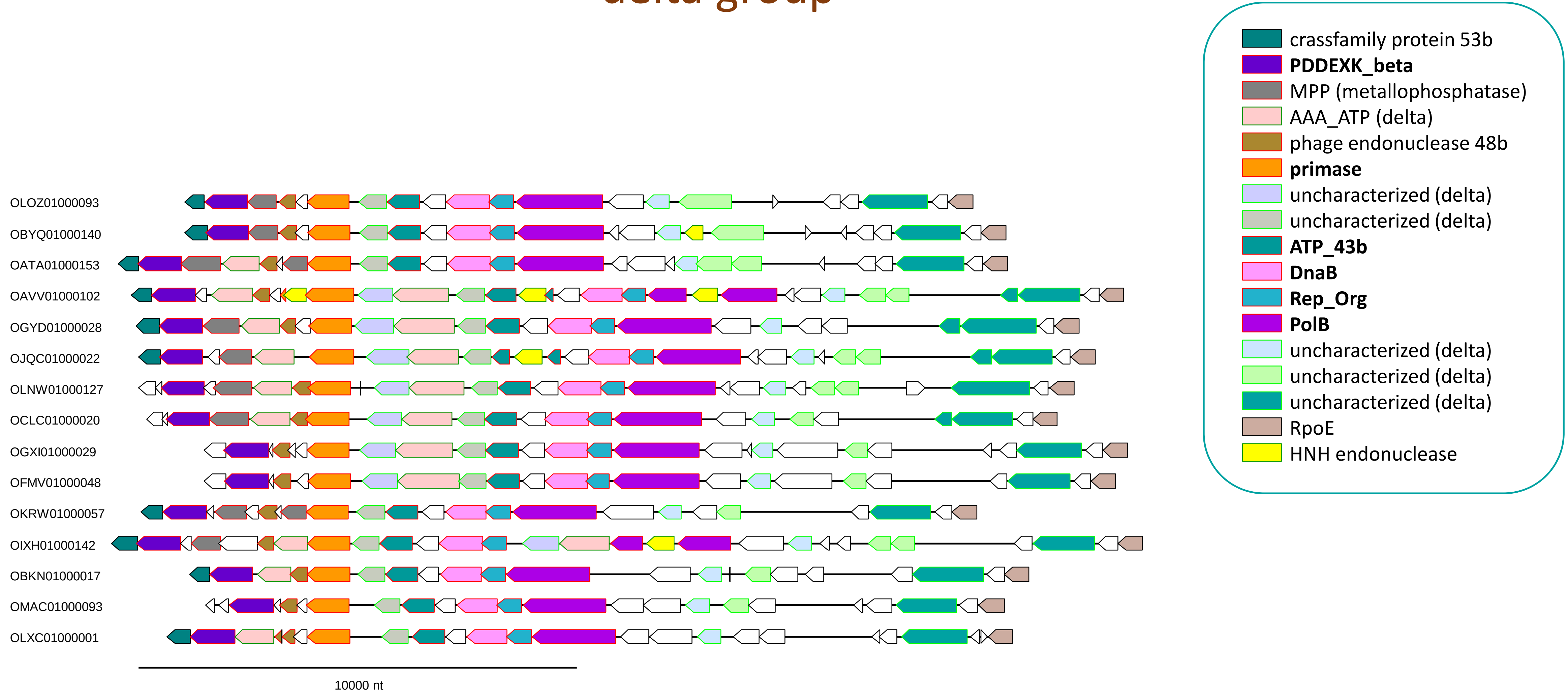

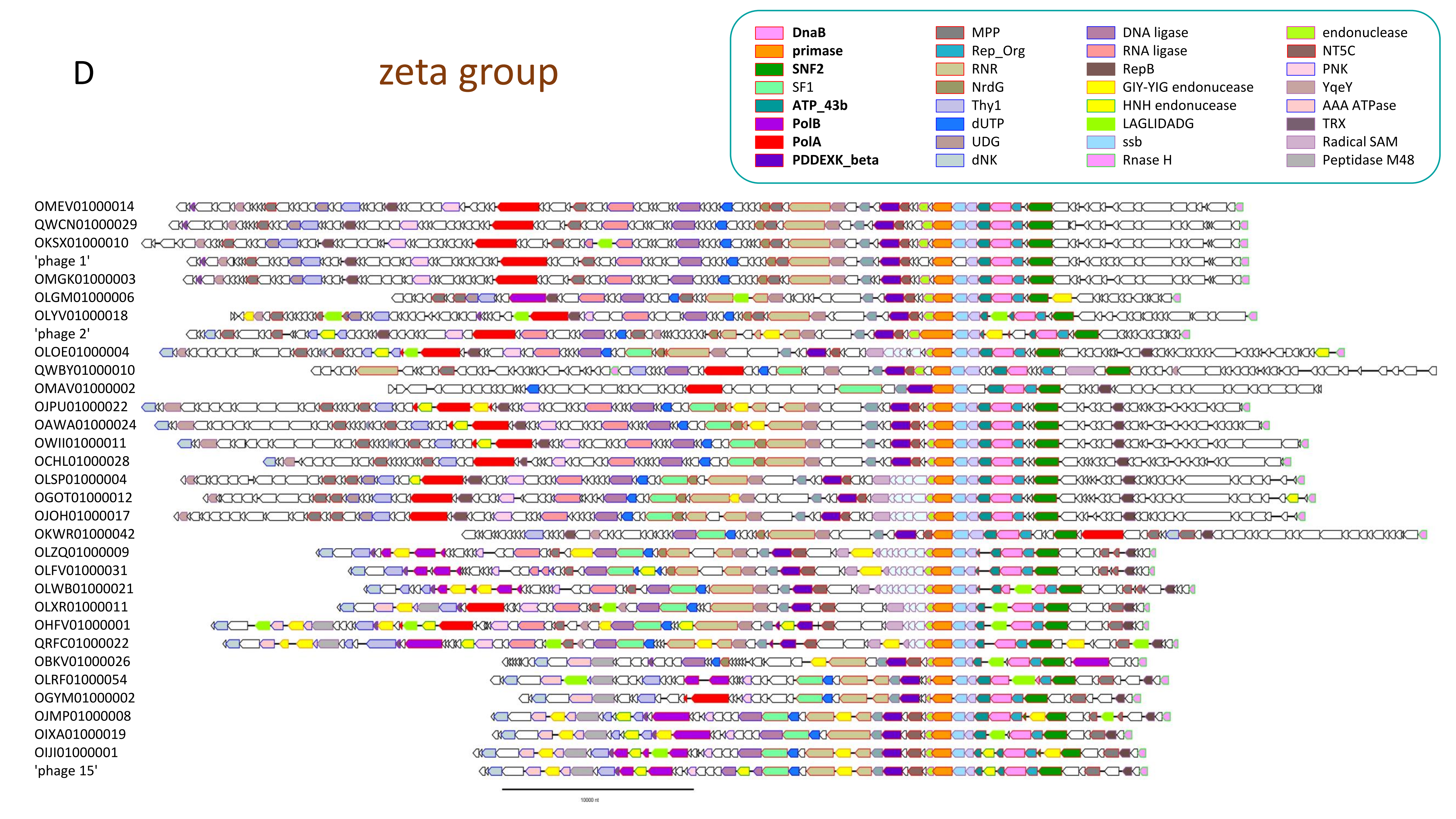
D
zeta group
DnaB
primase
SNF2
SF1
ATP_43b
PolB
PolA
PDDEXK_beta
MPP
Rep_Org
RNR
NrdG
Thy1
dUTP
UDG
dNK
DNA ligase
RNA ligase
RepB
GIY-YIG endonucease
HNH endonucease
LAGLIDADG
ssb
Rnase H
endonuclease
NT5C
PNK
YqeY
AAA ATPase
TRX
Radical SAM
Peptidase M48
OMEV01000014
QWCN01000029
OKSX01000010
'phage 1'
OMGK01000003
OLGM01000006
OLYV01000018
'phage 2'
OLOE01000004
QWBY01000010
OMAV01000002
OJPU01000022
OAWA01000024
OWII01000011
OCHL01000028
OLSP01000004
OGOT01000012
OJOH01000017
OKWR01000042
OLZQ01000009
OLFV01000031
OLWB01000021
OLXR01000011
OHFV01000001
QRFC01000022
OBKV01000026
OLRF01000054
OGYM01000002
OJMP01000008
OIXA01000019
OIJI01000001
'phage 15'
10000 nt

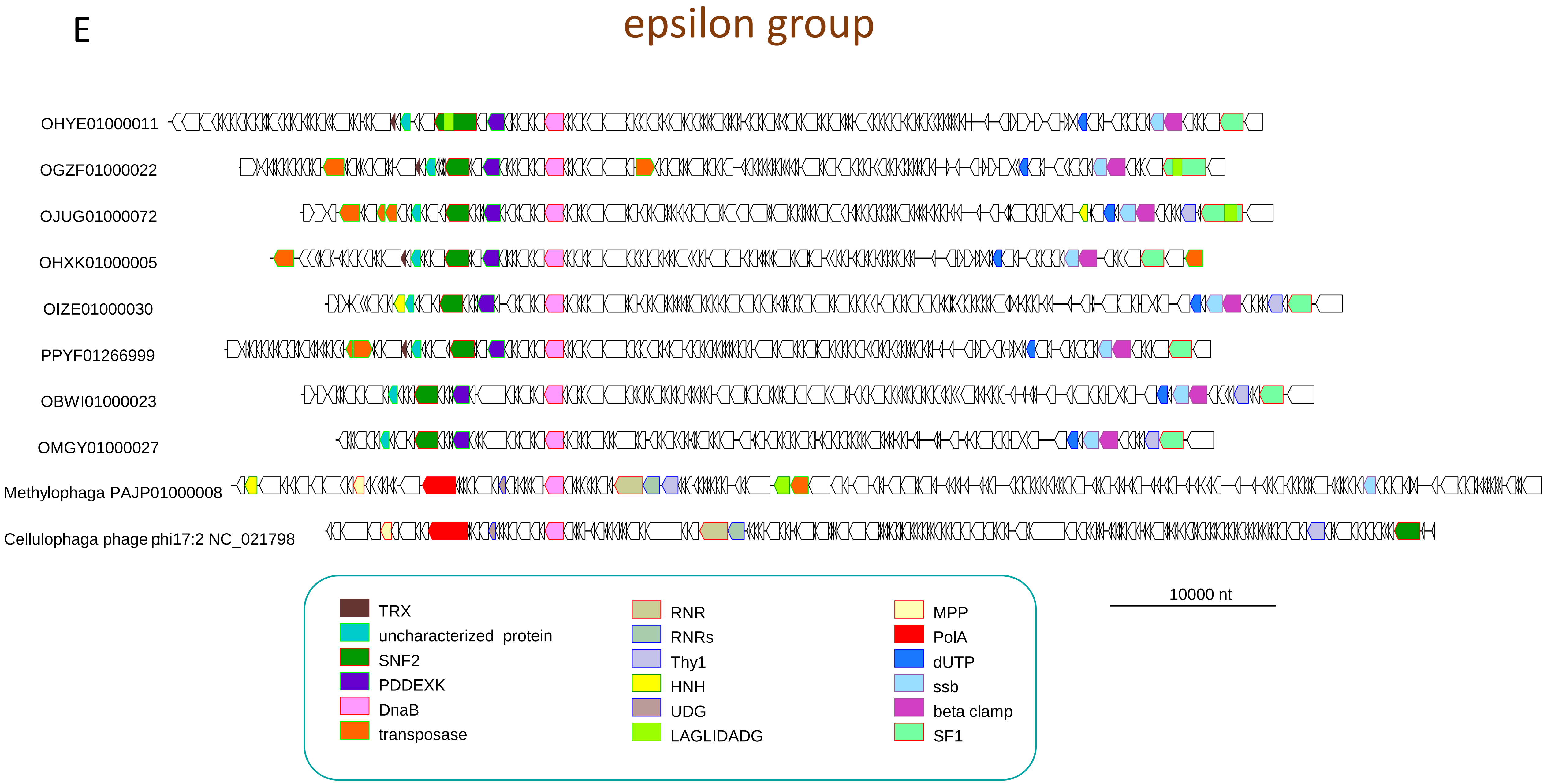
E
epsilon group
OHYE01000011
OGZF01000022
OJUG01000072
OHXK01000005
OIZE01000030
PPYF01266999
OBWI01000023
OMGY01000027
Methylophaga PAJP01000008
Cellulophaga phage phi17:2 NC_021798
TRX
uncharacterized protein
SNF2
PDDEXK
DnaB
transposase
RNR
RNRs
Thy1
HNH
UDG
LAGLIDADG
MPP
PolA
dUTP
ssb
beta clamp
SF1
10000 nt

### Supplementary Figure 2

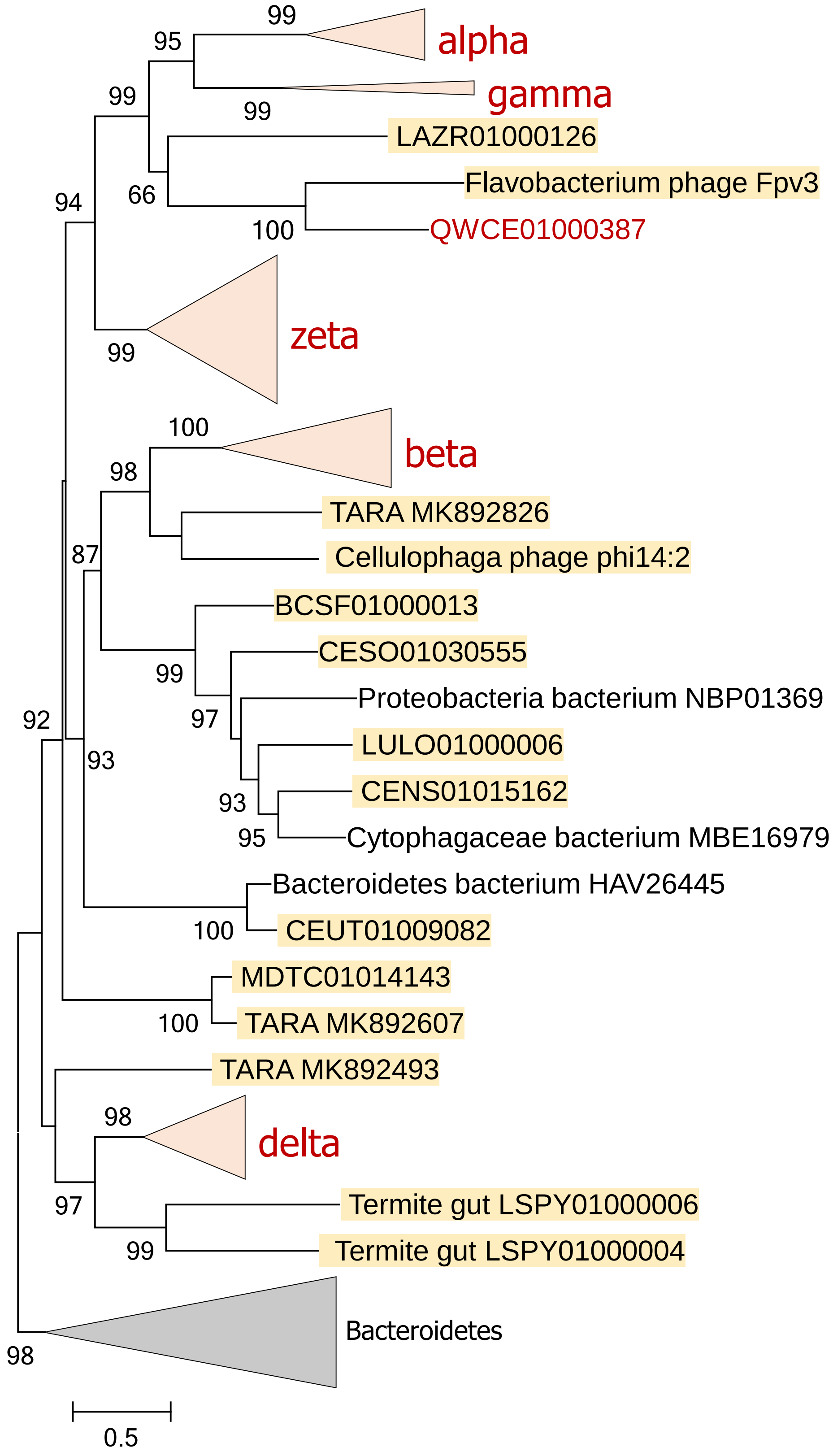

### Supplementary Figure 3

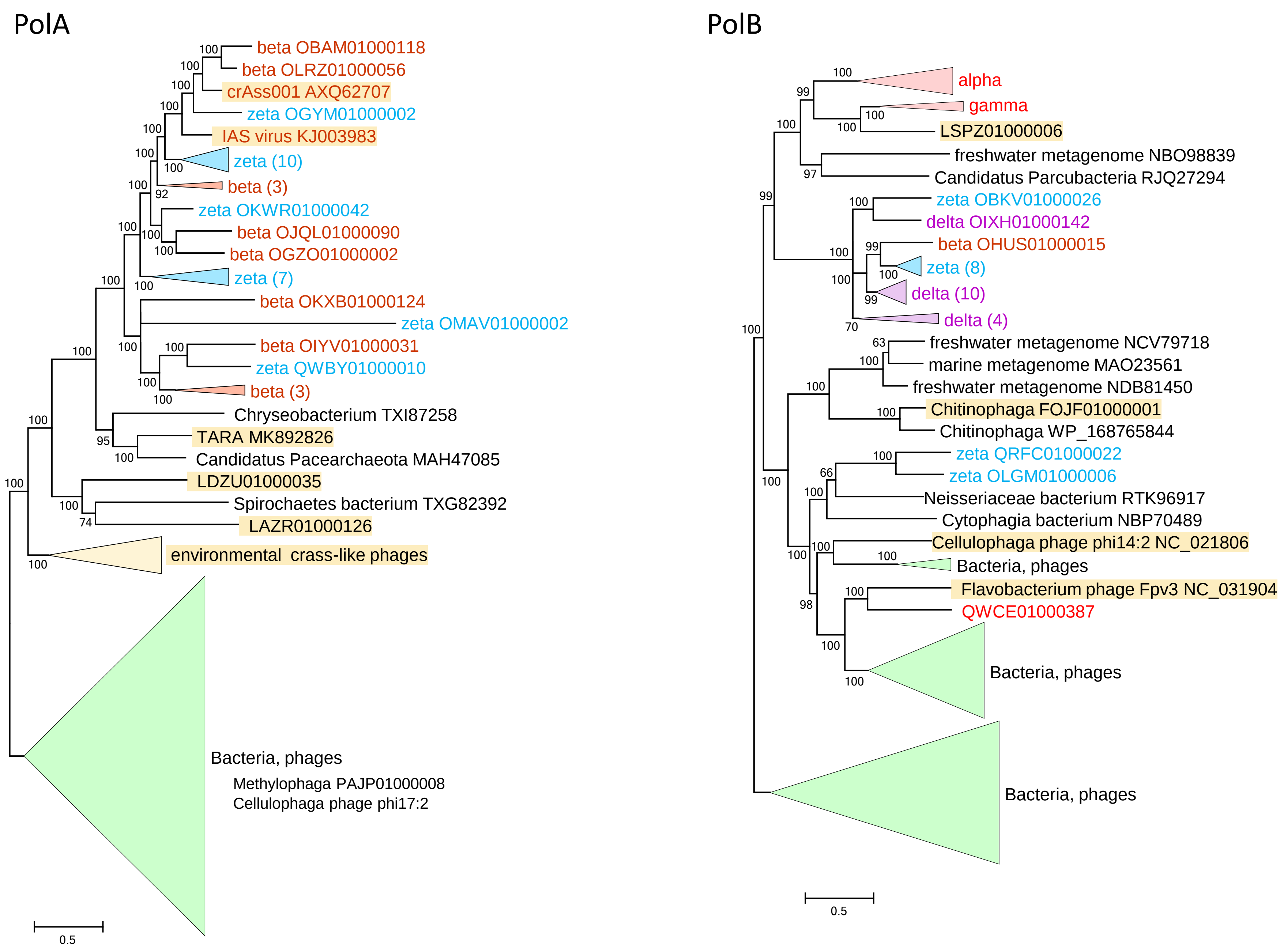

Supplementary Figure 4

A

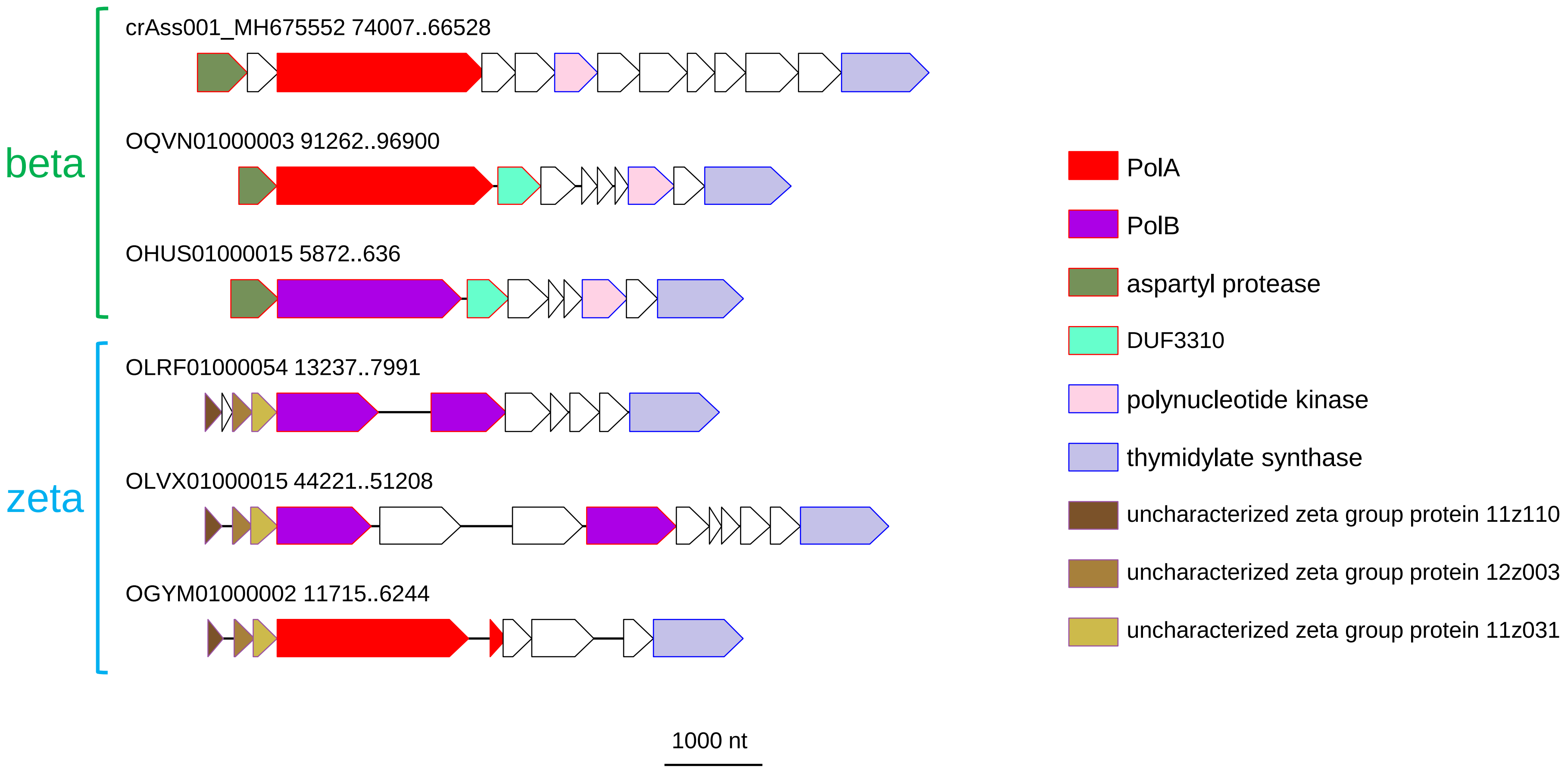

B

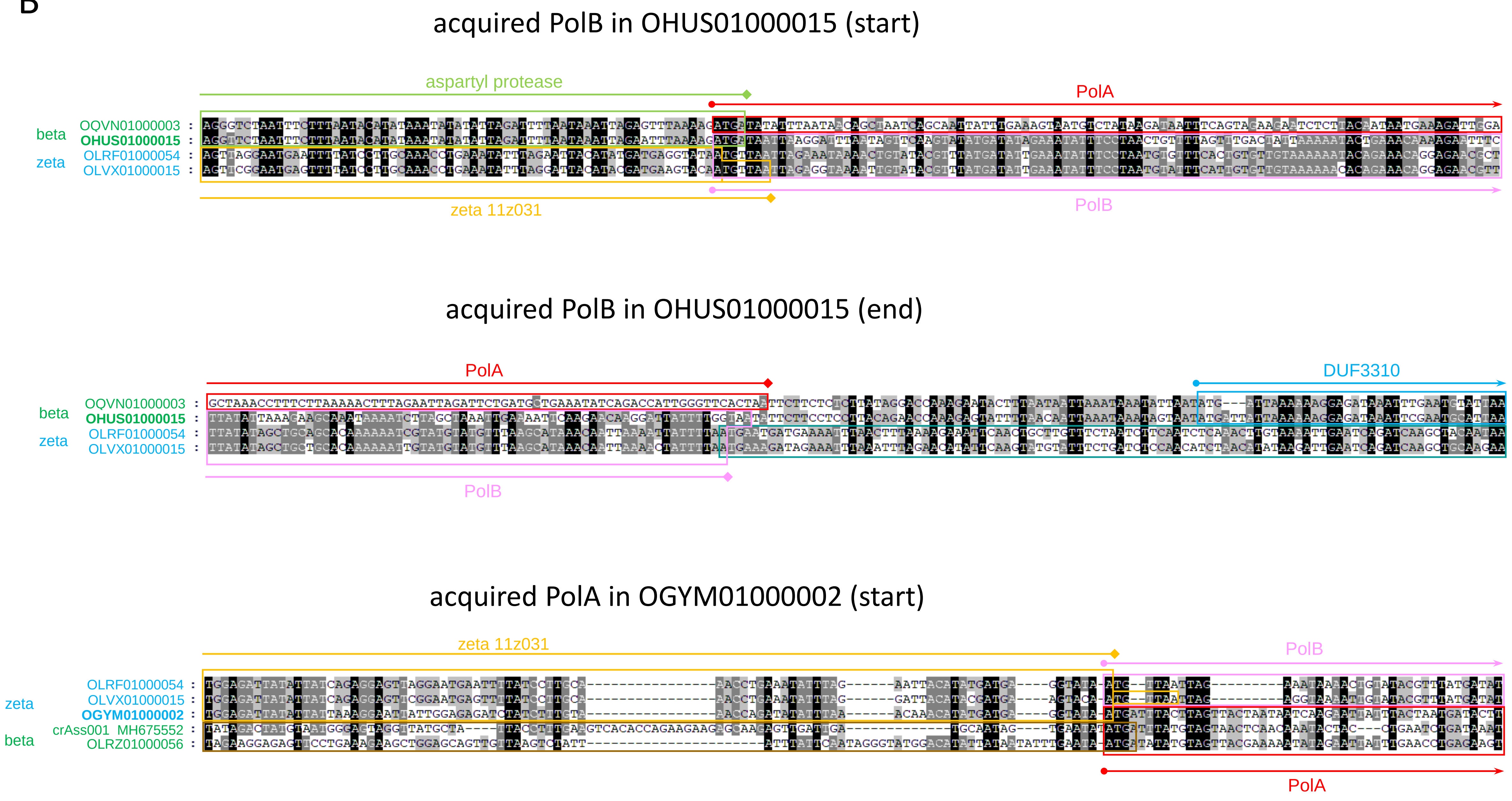
acquired PolB in OHUS01000015 (start)
aspartyl protease
PolA
beta
OQVN01000003
OHUS01000015
zeta
OLRF01000054
OLVX01000015
zeta 11z031
PolB
acquired PolB in OHUS01000015 (end)
PolA
DUF3310
beta
OQVN01000003
OHUS01000015
zeta
OLRF01000054
OLVX01000015
PolB
acquired PolA in OGYM01000002 (start)
zeta 11z031
PolB
zeta
OLRF01000054
OLVX01000015
OGYM01000002
beta
crAss001 MH675552
OLRZ01000056
PolA

### Supplementary Figure 5

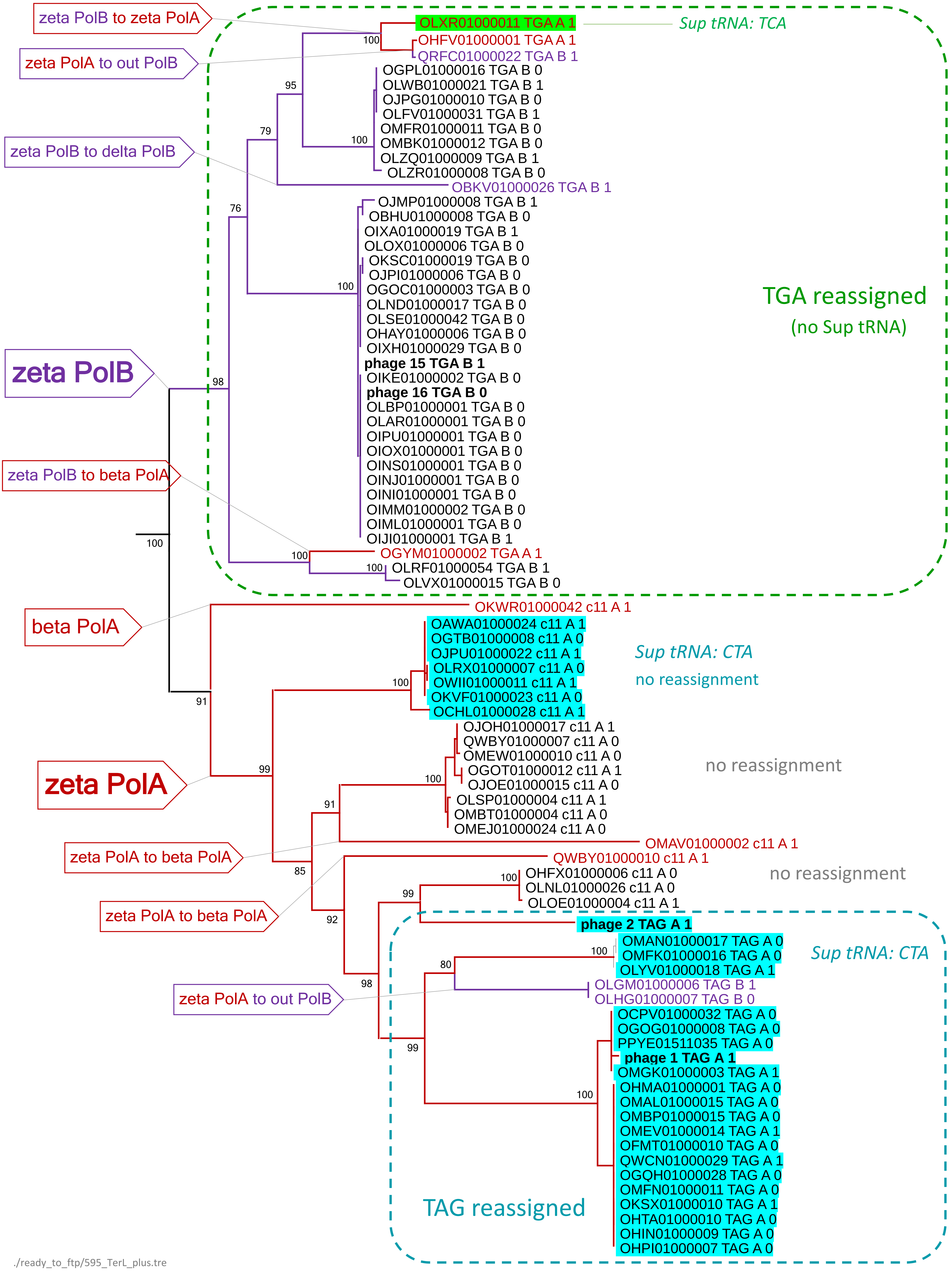


./ready_to_ftp/595_TerL_plus.tre

### Supplementary Figure 6

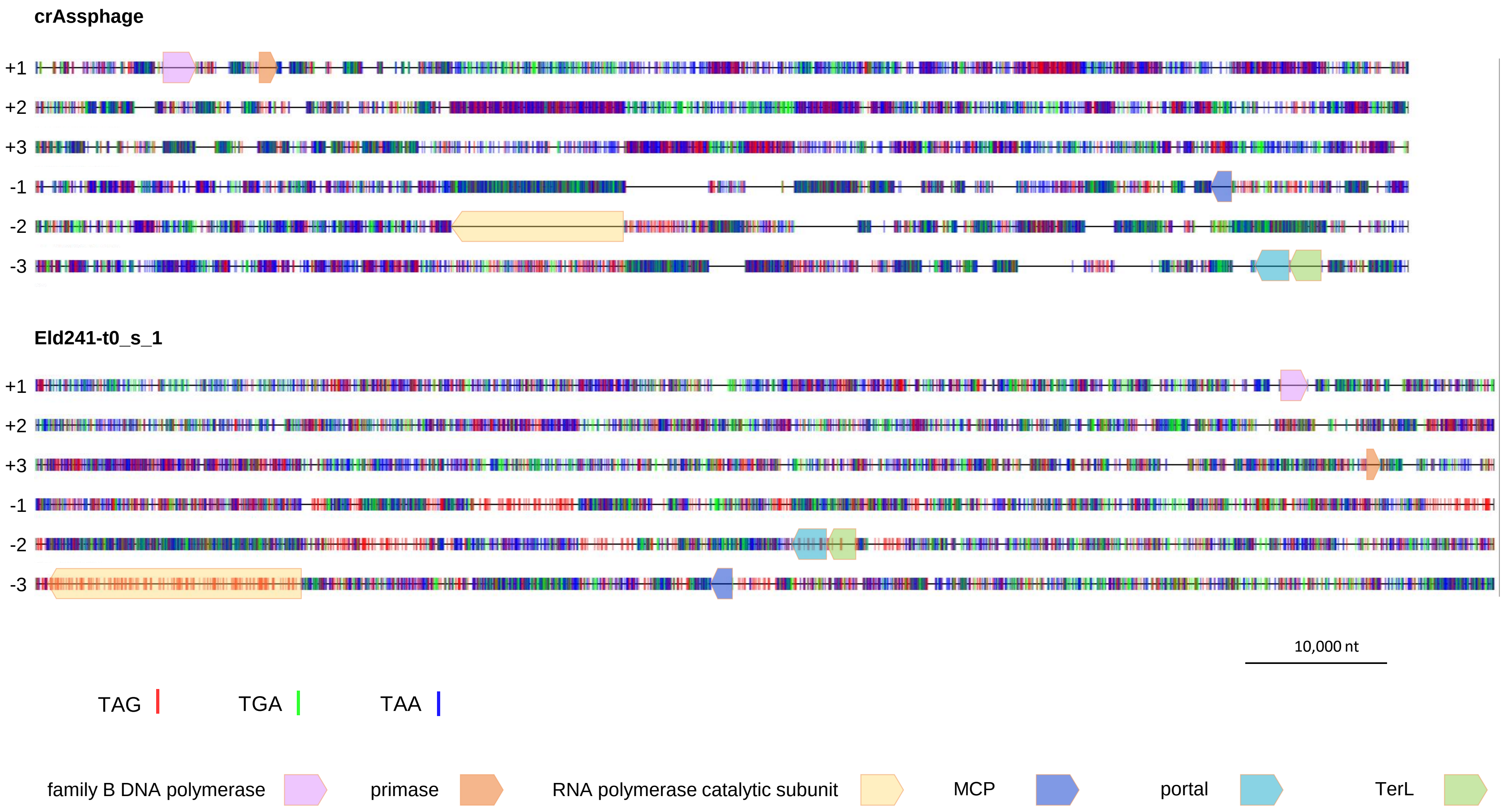

Supplementary Figure 7

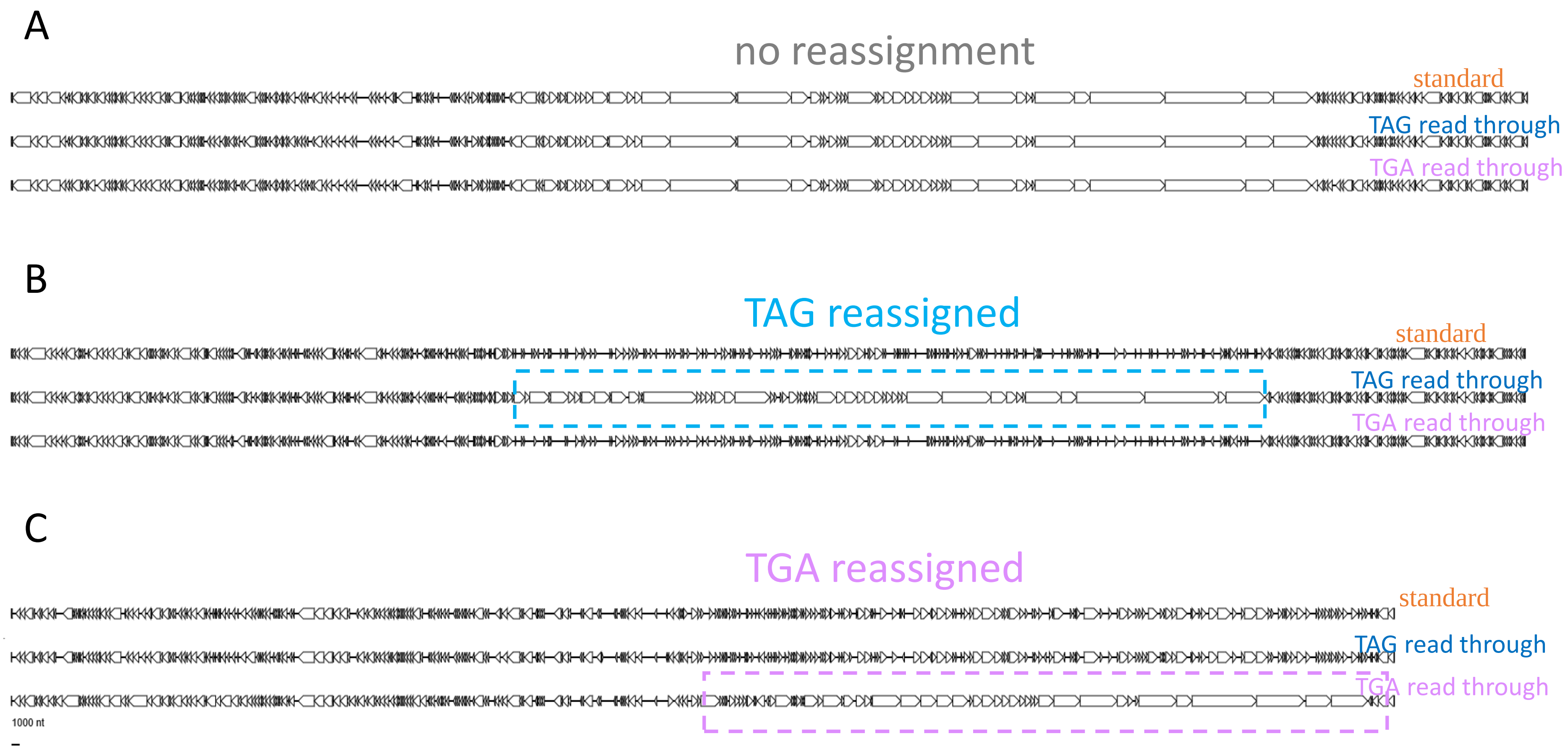

### Supplementary Figure 8

A

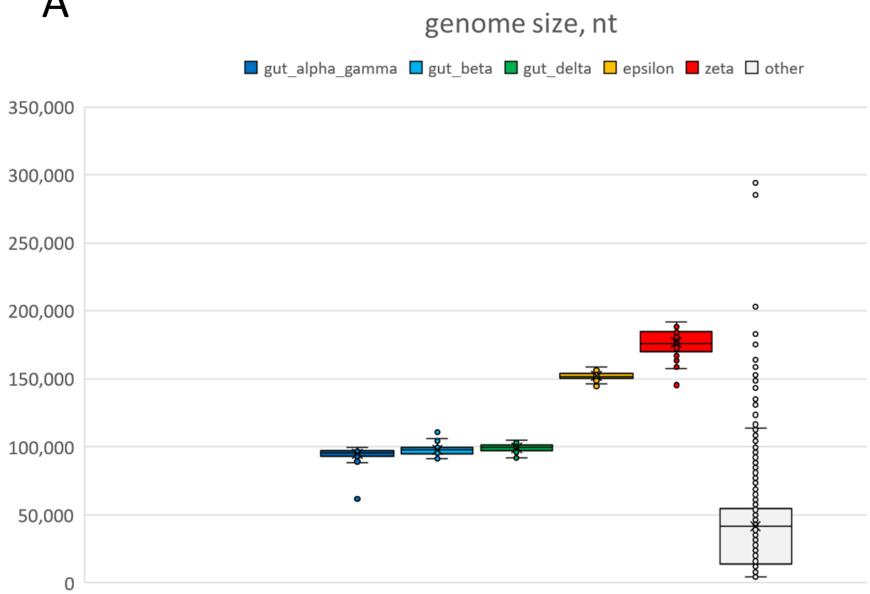


B

| | # of contigs | # of tRNA | # of Sup tRNA | anticodon | reassigned codons |
|---|---|---|---|---|---|
| alpha gamma | 190 | 115 | 1 | TTA | TAA |
| beta | 57 | 595 | 1 | CTA | TAG |
| delta | 233 | 472 | 153 | CTA | TAG |
| epsilon | 36 | 132 | 0 | | |
| zeta | 79 | 1662 | 27 | 26 CTA, 1 TCA | 26 TAG, 1 TCA |
| Flavob_phages | 1 | 10 | 0 | | |
| other cMAGs | 3343 | 2203 | 9 | 1 TCA, 2 CTA, 6 TTA | 1 TCA, 2 CTA, 6 TTA |

C

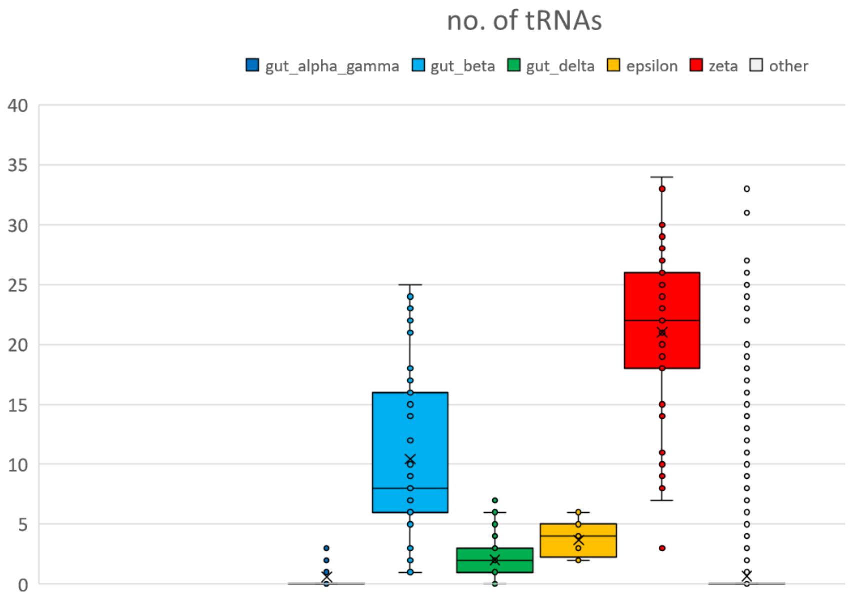


D

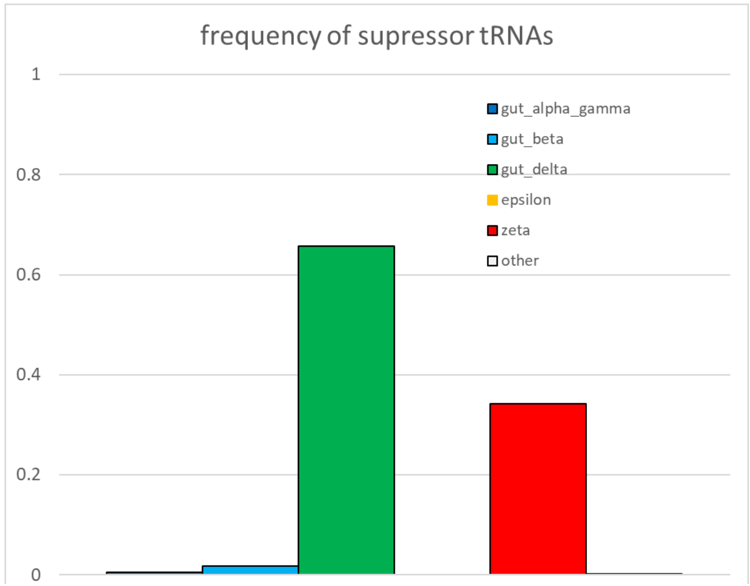

Supplementary Figure 9

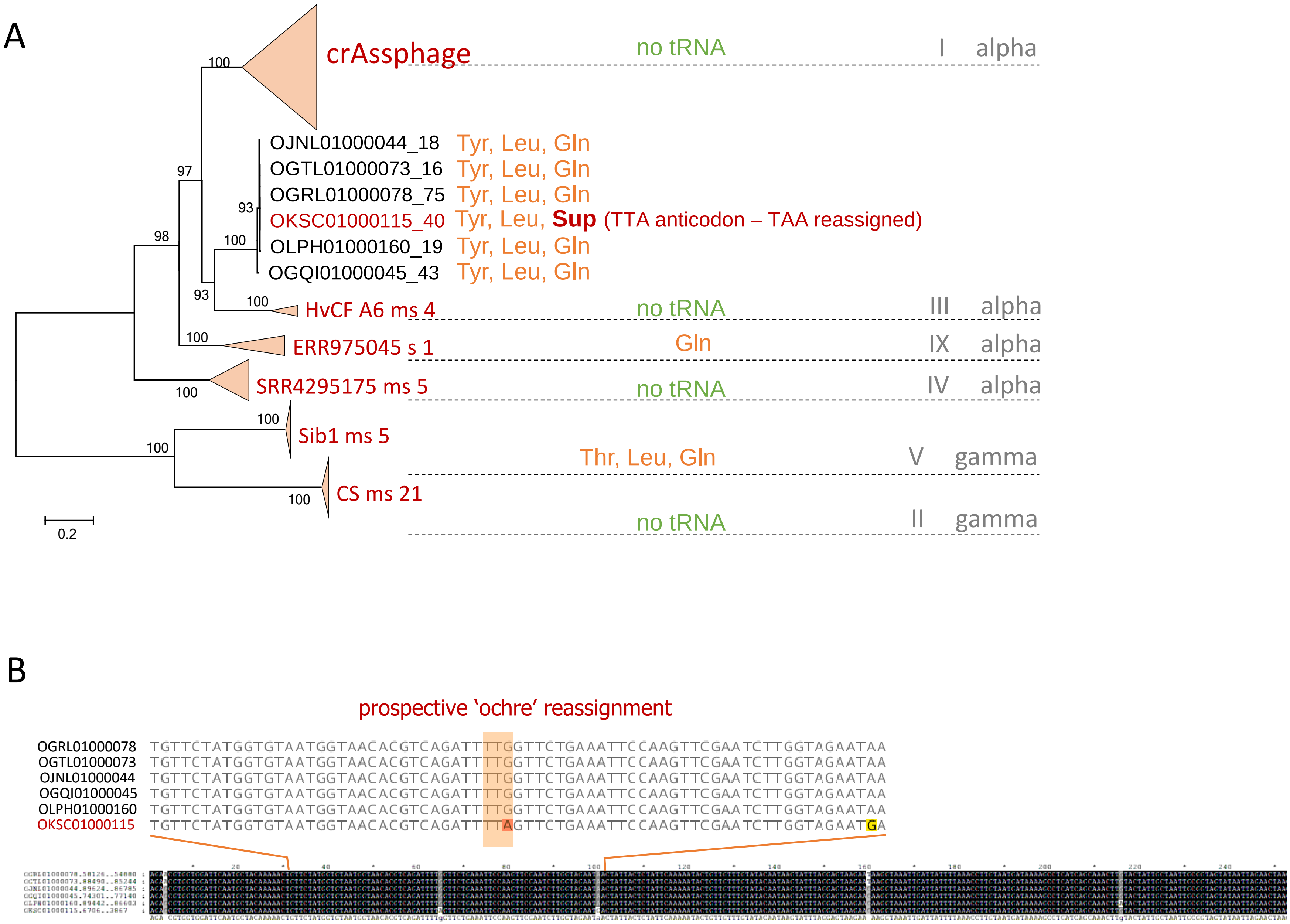

Supplementary Figure 10

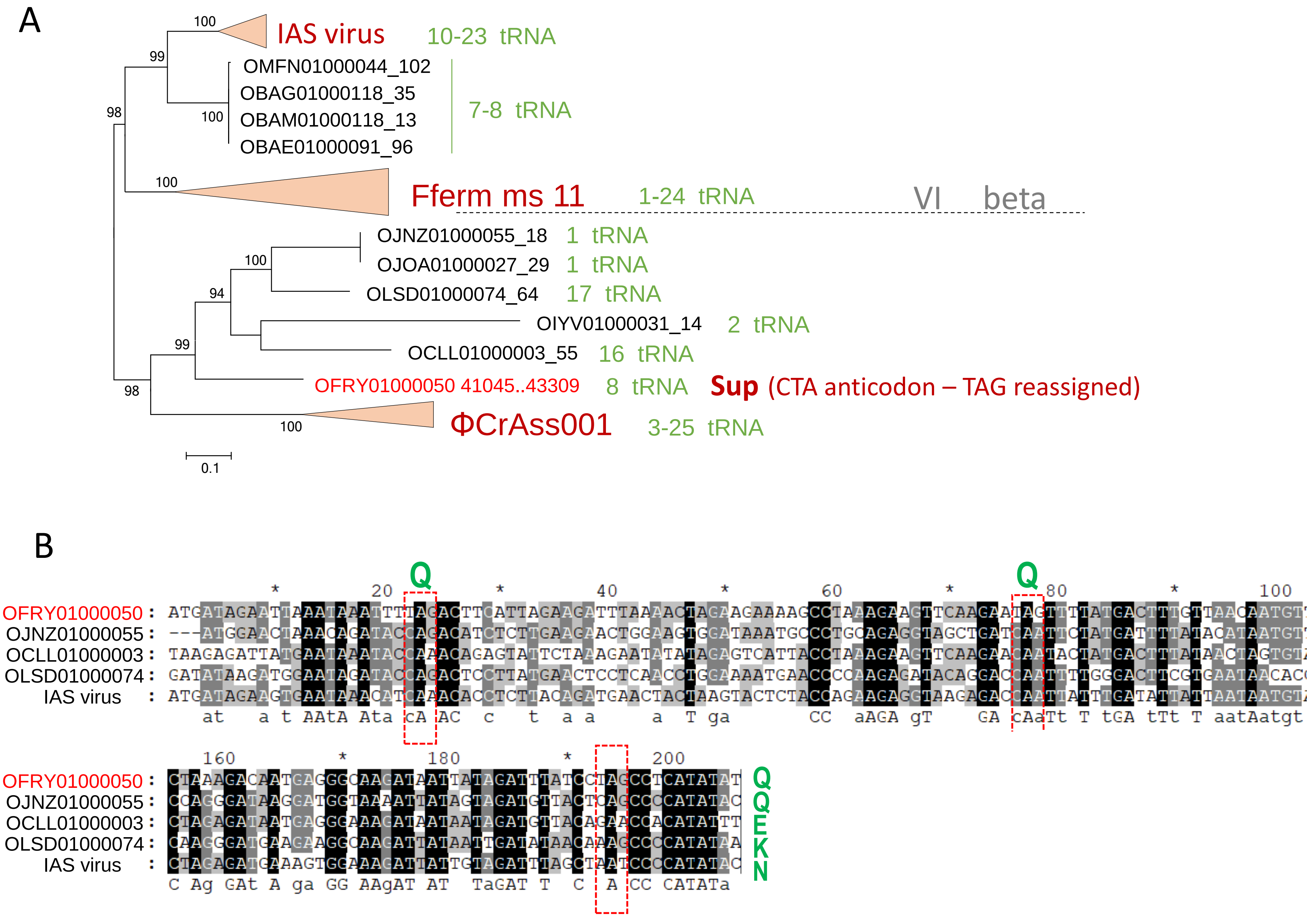

C

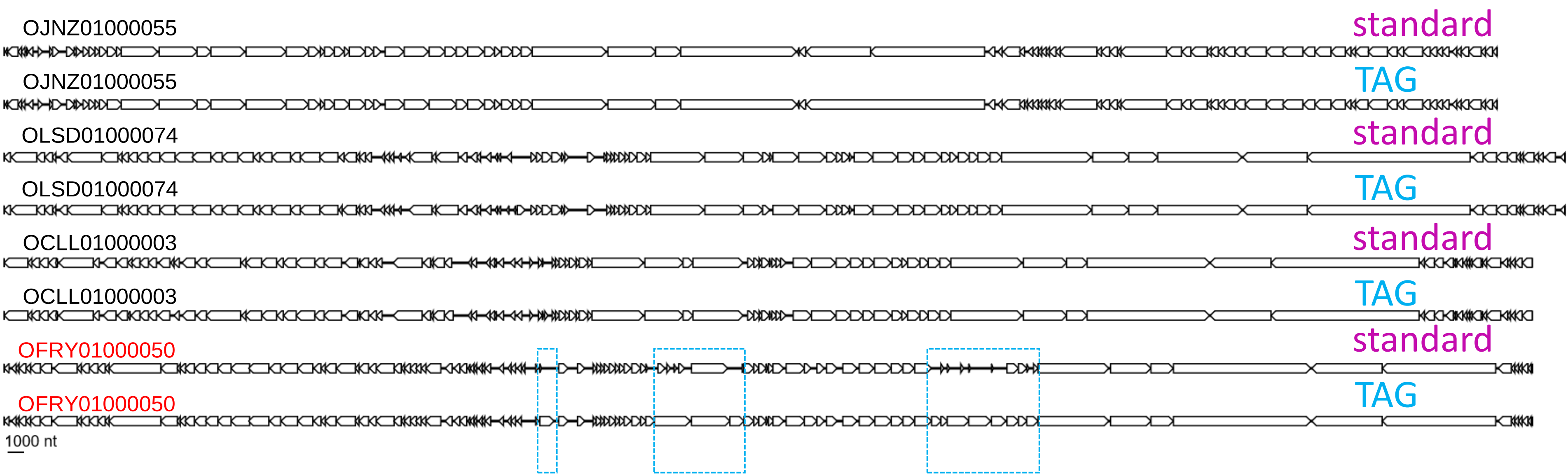

OJNZ01000055
standard
OJNZ01000055
TAG
OLSD01000074
standard
OLSD01000074
TAG
OCLL01000003
standard
OCLL01000003
TAG
OFRY01000050
standard
OFRY01000050
TAG
1000 nt

### Supplementary Figure 11

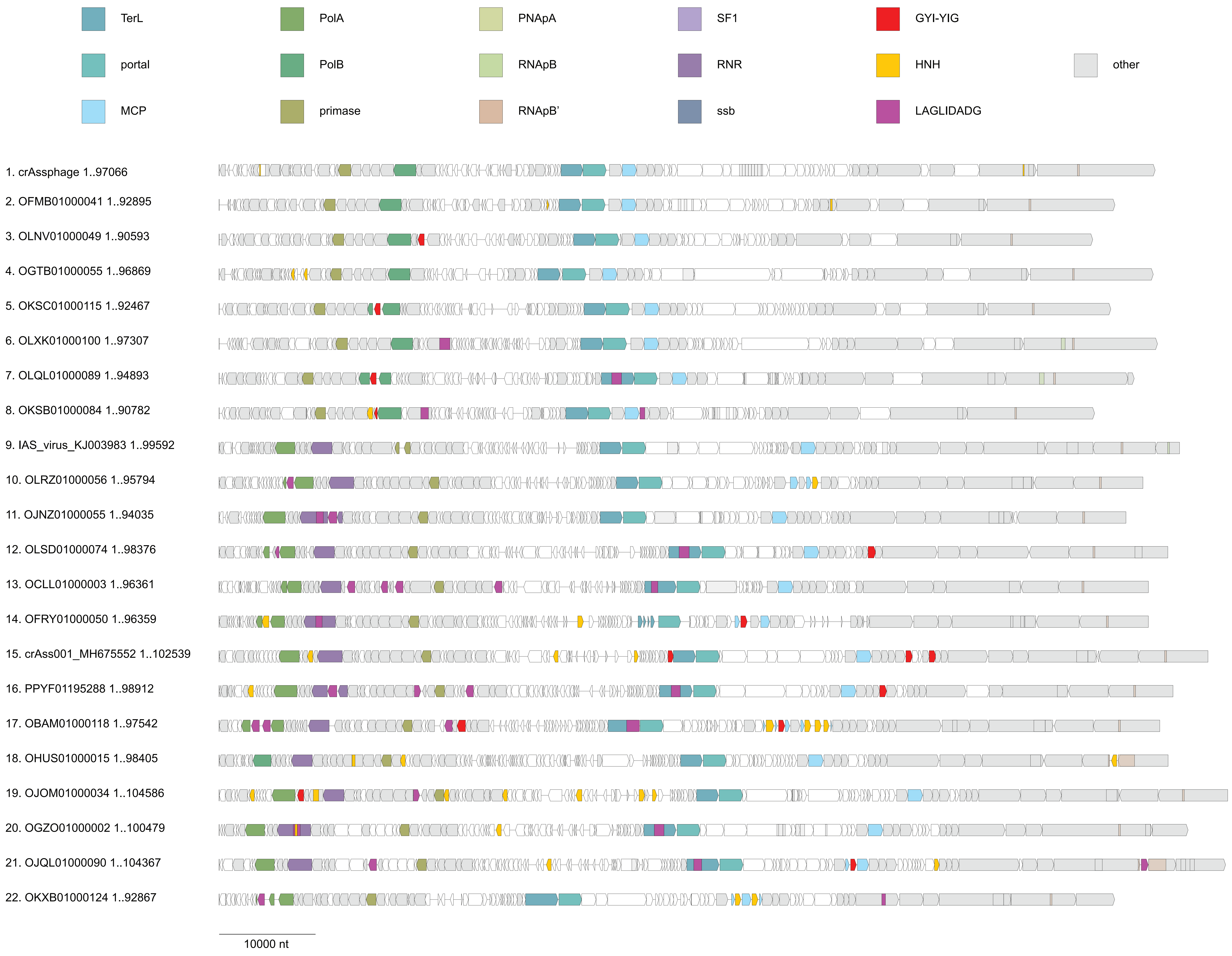

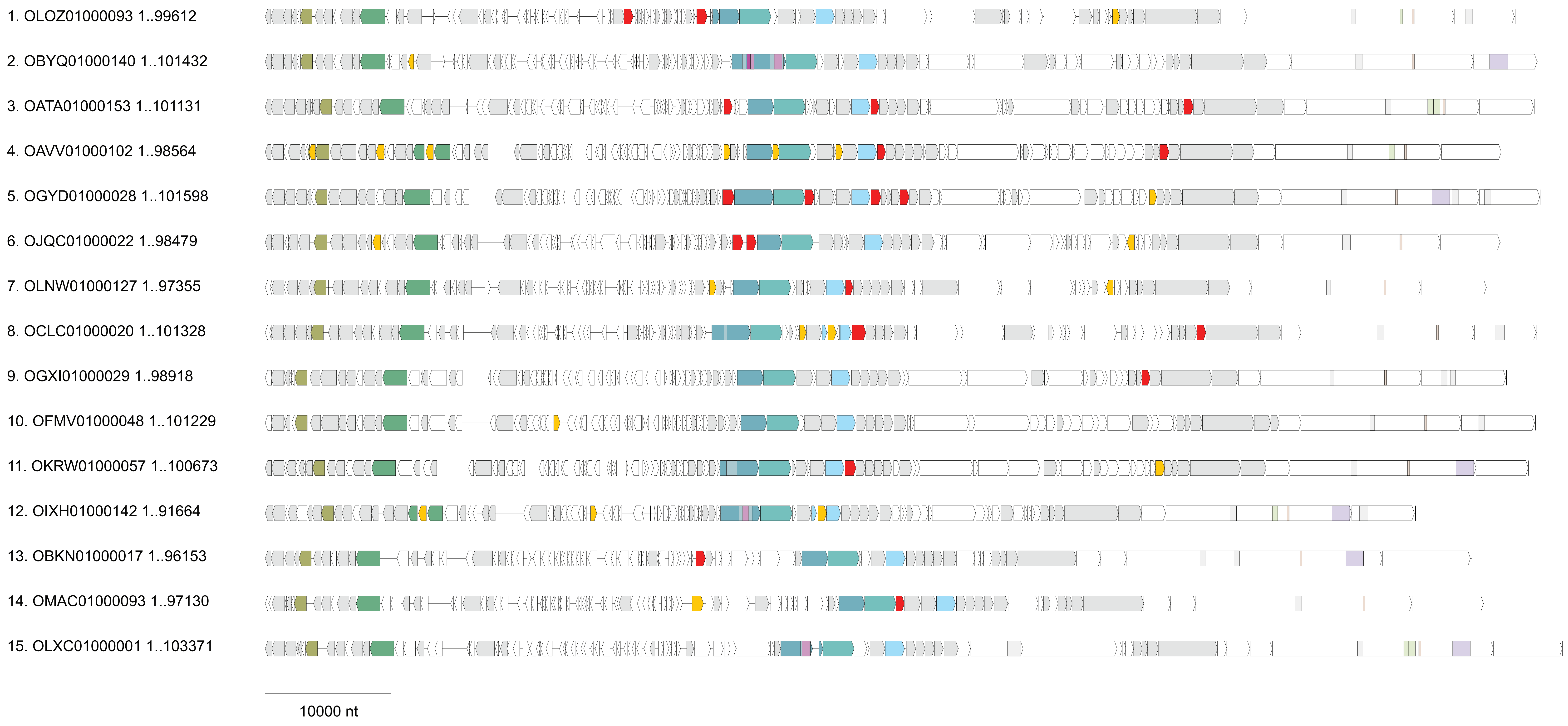
1. OLOZ01000093 1..99612
2. OBYQ01000140 1..101432
3. OATA01000153 1..101131
4. OAVV01000102 1..98564
5. OGYD01000028 1..101598
6. OJQC01000022 1..98479
7. OLNW01000127 1..97355
8. OCLC01000020 1..101328
9. OGXI01000029 1..98918
10. OFMV01000048 1..101229
11. OKRW01000057 1..100673
12. OIXH01000142 1..91664
13. OBKN01000017 1..96153
14. OMAC01000093 1..97130
15. OLXC01000001 1..103371
10000 nt

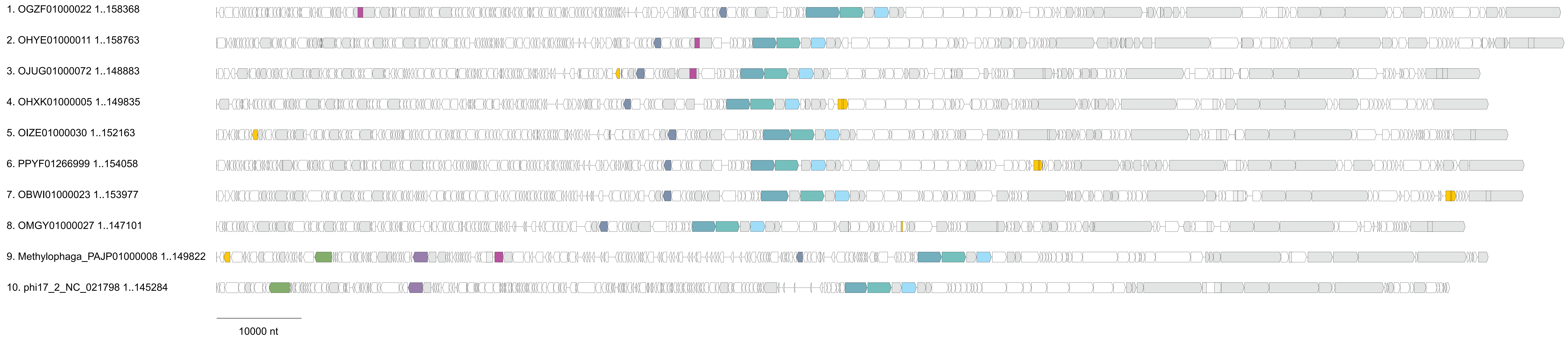
1. OGZF01000022 1..158368
2. OHYE01000011 1..158763
3. OJUG01000072 1..148883
4. OHXK01000005 1..149835
5. OIZE01000030 1..152163
6. PPYF01266999 1..154058
7. OBWI01000023 1..153977
8. OMGY01000027 1..147101
9. Methylophaga_PAJP01000008 1..149822
10. phi17_2_NC_021798 1..145284
10000 nt

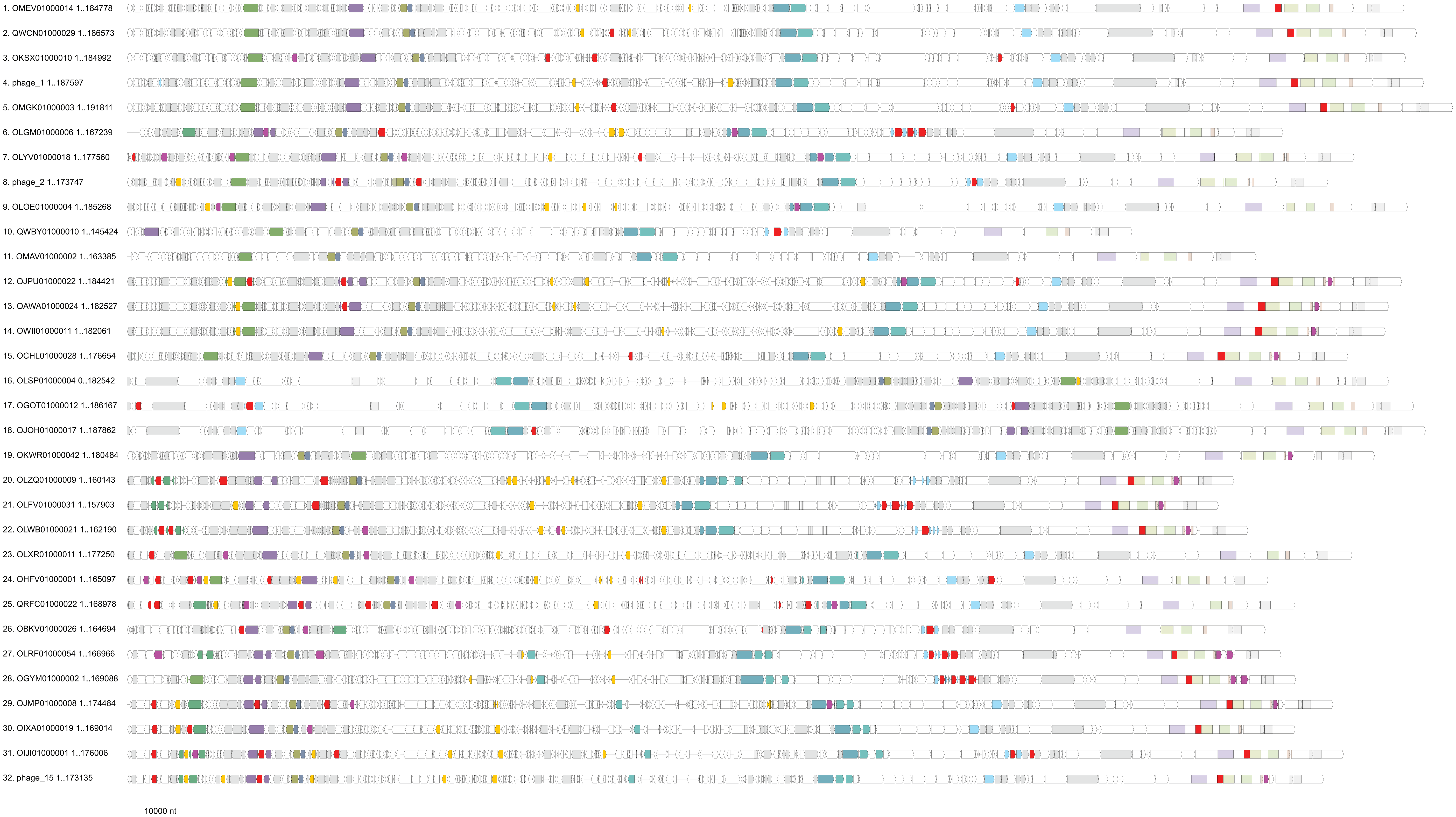
1. OMEV01000014 1..184778
2. QWCN01000029 1..186573
3. OKSX01000010 1..184992
4. phage_1 1..187597
5. OMGK01000003 1..191811
6. OLGM01000006 1..167239
7. OLYV01000018 1..177560
8. phage_2 1..173747
9. OLOE01000004 1..185268
10. QWBY01000010 1..145424
11. OMAV01000002 1..163385
12. OJPU01000022 1..184421
13. OAWA01000024 1..182527
14. OWII01000011 1..182061
15. OCHL01000028 1..176654
16. OLSP01000004 0..182542
17. OGOT01000012 1..186167
18. OJOH01000017 1..187862
19. OKWR01000042 1..180484
20. OLZQ01000009 1..160143
21. OLFV01000031 1..157903
22. OLWB01000021 1..162190
23. OLXR01000011 1..177250
24. OHFV01000001 1..165097
25. QRFC01000022 1..168978
26. OBKV01000026 1..164694
27. OLRF01000054 1..166966
28. OGYM01000002 1..169088
29. OJMP01000008 1..174484
30. OIXA01000019 1..169014
31. OIJI01000001 1..176006
32. phage_15 1..173135
10000 nt

### Supplementary Figure 12

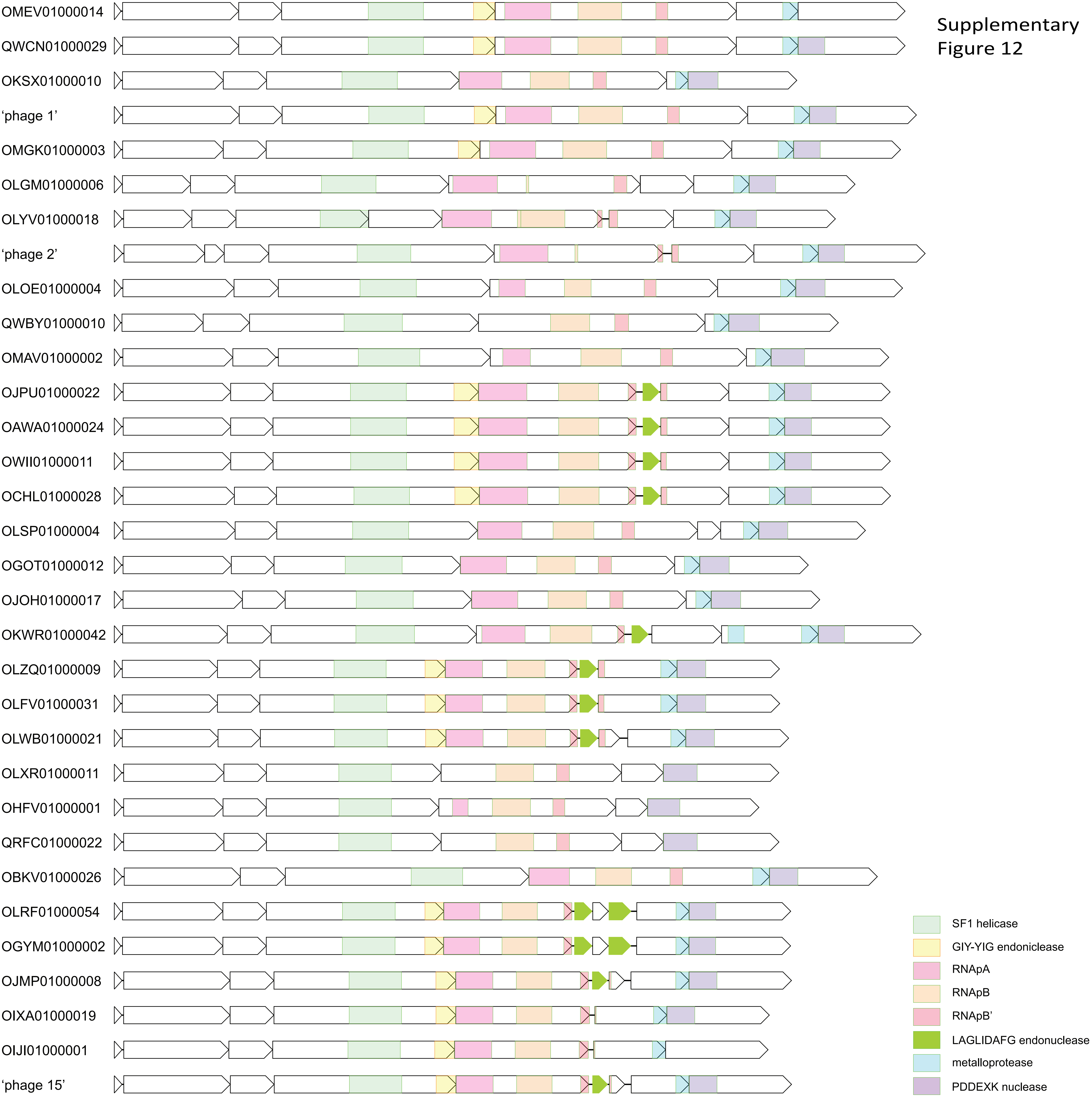
